# *Plasmodium* Kinesin-8X associates with mitotic spindles and is essential for oocyst development during parasite proliferation and transmission

**DOI:** 10.1101/665836

**Authors:** Mohammad Zeeshan, Fiona Shilliday, Tianyang Liu, Steven Abel, Tobias Mourier, David J. P. Ferguson, Edward Rea, Rebecca R. Stanway, Magali Roques, Desiree Williams, Emilie Daniel, Declan Brady, Anthony J. Roberts, Anthony A. Holder, Arnab Pain, Karine G. Le Roch, Carolyn A. Moores, Rita Tewari

**Affiliations:** School of Life Sciences, Queens Medical Centre, University of Nottingham, Nottingham, NG7 2UH, UK; Institute of Structural and Molecular Biology, Department of Biological Sciences, Birkbeck College, London, WC1E 7HX, United Kingdom; Department of Molecular, Cell and Systems Biology, University of California Riverside, 900 University Ave, Riverside, CA, 92521, USA; Computational Bioscience Research Center, Biological Environmental Sciences and Engineering Division, King Abdullah University of Science and Technology, Thuwal, Jeddah 23955-6900, Kingdom of Saudi Arabia; Nuffield Department of Clinical Laboratory Science, University of Oxford, John Radcliffe Hospital, Oxford OX3 9DU, UK; Department of Biological and Medical Sciences, Faculty of Health and Life Science, Oxford Brookes University, Gipsy Lane, Oxford OX3 0BP, UK; Institute of Cell Biology, University of Bern, Bern 3012, Switzerland; Malaria Parasitology Laboratory, The Francis Crick Institute, London, NW1 1AT, UK

## Abstract

Kinesin-8 proteins are microtubule motors that are often involved in regulation of mitotic spindle length and chromosome alignment. They move towards the ends of spindle microtubules and regulate the dynamics of these ends due, at least in some species, to their microtubule depolymerization activity. *Plasmodium spp*. exhibit an atypical endomitotic cell division in which chromosome condensation and spindle dynamics are not well understood in the different proliferative stages. Genome-wide homology analysis of *Plasmodium spp*. revealed the presence of two Kinesin-8 motor proteins (Kinesin-8X and Kinesin-8B). Here we have studied the biochemical properties of Kinesin-8X and its role in parasite proliferation. *In vitro*, Kinesin-8X showed motile and depolymerization activities like other Kinesin-8 motors. To understand its role in cell division, we have used protein tagging and live cell imaging to define the location of *Plasmodium* Kinesin-8X during all proliferative stages of the *P berghei* life cycle. Furthermore, we have used gene targeting to analyse the function of Kinesin-8X. The results reveal a spatio-temporal involvement of Kinesin-8X in spindle dynamics and its association with both mitotic and meiotic spindles and the putative microtubule organising centre (MTOC). Deletion of the Kinesin-8X gene showed that this protein is required for endomitotic division during oocyst development and is therefore necessary for parasite replication within the mosquito gut, and for transmission to the vertebrate host. Consistently, transcriptome analysis of *Δkinesin-8X* parasites reveals modulated expression of genes involved mainly in microtubule-based processes, chromosome organisation and the regulation of gene expression supporting a role in cell division.

**Author Summary:** Kinesins are microtubule-based motors that play key roles in intracellular transport, cell division and motility. Members of the Kinesin-8 family contribute to chromosome alignment during cell division in many eukaryotes. However, the roles of kinesins in the atypical cell division in *Plasmodium*, the causative agent of malaria, is not known. In contrast to many other eukaryotes, *Plasmodium* proliferates by endomitosis, in which genome replication and division occur within a nucleus bounded by a persistent nuclear envelope. We show that the *Plasmodium* genome encodes only nine kinesins and we further investigate the role of Kinesin-8X throughout the *Plasmodium* life cycle using biochemical and gene targeting approaches. We show that *Plasmodium* Kinesin-8X has microtubule-based motility and depolymerization activity. We also show that Kinesin-8X is probably localized on putative MTOCs and spindles during cell division in most of the stages of *P. berghei* life cycle. By gene deletion we demonstrate that Kinesin-8X is essential for normal oocyst development and sporozoite formation. Genome-wide RNA analysis of *Δkinesin-8X* parasites reveals modulated expression of genes involved in microtubule-based processes. Overall, the data suggest that Kinesin-8X is a molecular motor that plays essential roles during endomitosis in oocyst development in the mosquito, contributing to parasite transmission.

## Introduction

Kinesins are molecular motors that use ATP to translocate along microtubules (MTs) or control MT-end dynamics. There are 14 to 16 classes of kinesins in eukaryotes [1–3] defined by their conserved motor domains, which are located in different contexts within the protein primary sequence, and which undertake distinct cellular functions [4]. Many kinesins together with dynein have important roles in mitosis, including spindle pole separation, kinetochore attachment to spindles, chromosome alignment and segregation and cytokinesis [5, 6]. Some of these kinesins have also been shown to play an essential role in meiosis, mainly during meiosis I, where recombination takes place. [7–9]. Kinesin-8 is one class of these motors that is conserved across eukaryotes [2, 3]. During mitosis, Kinesin-8 proteins in many eukaryotes localise to spindles and control spindle length and chromosome positioning at the cell equator [10–14]. In the absence of functional Kinesin-8, the mitotic spindle length increases and chromosome alignment at the metaphase plate is affected [10, 15, 16]. At the molecular level, they are plus end directed MT motors that play a key role in controlling MT length, and some exhibit MT depolymerisation activity [17–19]. In addition, Kinesin-8 proteins may have a role in maintenance of cell polarity and nuclear positioning in the centre of a fission yeast [20–22].

Malaria is the most deadly parasitic disease, and is caused by the unicellular protozoan *Plasmodium* spp. belonging to subclass in the Apicomplexa phylum, which infects many vertebrates and is transmitted by female *Anopheles* mosquitoes [23]. The parasite has a complex life cycle, alternating between its two hosts. Proliferation is by closed endomitotic division in which genome replication and division occur within a nucleus bounded by a persistent nuclear envelope [24, 25]. Both the replication rate and the number of rounds of division varies between different *Plasmodium* species, at different stages of the life cycle, and within different hosts and host cells [24, 25]. During asexual stages, nuclear division is asynchronous and followed by synchronous cytokinesis to produce multiple haploid progeny, and is called schizogony within vertebrate hepatocytes and erythrocytes, and sporogony within oocysts attached to the mosquito gut basal lamina [25, 26]. Haploid sexual progenitor cells, male and female gametocytes, remain arrested early in the cell cycle within red blood cells and produce gametes only following ingestion in a blood meal within a mosquito gut where environmental conditions, including temperature, pH and mosquito factors such as xanthurenic acid, are optimal for gametocyte activation [27, 28]. Male gametocytes undergo three successive rounds of rapid DNA replication producing an 8N nucleus within 15 minutes of activation, followed by exflagellation to release eight flagellated male gametes [29, 30]. *Plasmodium* lacks a classical centriole to nucleate spindle microtubules and drive spindle formation; however there is evidence from electron microscopy for the presence of a putative microtubule organizing centre (MTOC) embedded in the nuclear membrane, which initiates the polymerization of spindle microtubules [31, 32]. This finding has been corroborated recently using live cell imaging with a tagged centrin, *Pb*CEN-4-GFP [33]. As in other eukaryotes, these spindle microtubules are attached to the sister chromatids and at the end of mitosis move towards the spindle poles to separate the chromosomes [31, 34]. This separation of chromosomes is aided by various proteins associated with spindle MTs, including the kinesin motors that regulate spindle dynamics and polymerization [35, 36]. However, the role of kinesins in spindle dynamics during the atypical cell division of *Plasmodium* has not been studied.

Phylogenetic analyses have identified 9 kinesin genes in the *Plasmodium* genome [2, 3, 37] though not much is known in other members of the group Apicomplexa. Here we have analysed a high-resolution representation of the phylogenetic distribution of kinesins in all known Apicomplexa including *Plasmodium*. Earlier studies and the present one have identified two kinesins that are classified as members of the Kinesin-8 family which, according to the classification scheme of Wickstead and colleagues [3], are members of distinct Kinesin-8 subgroups, Kinesin-8B and Kinesin-8X. However, little is known about their function, location or dynamics in spindle formation during chromosome separation and atypical nuclear division in the parasite. To understand the role of *Plasmodium* Kinesin-8X we analysed its biochemical properties, focusing on both *P. falciparum* and *P. berghei* proteins, and its location and function throughout the entire life cycle using the rodent malaria parasite, *P. berghei*. We demonstrate that the motor domain is a MT-stimulated ATPase that drives MT gliding and has MT depolymerization activity. These activities are conserved in both *P. berghei* and *P. falciparum* Kinesin-8X. Live cell imaging of *P. berghei* showed that Kinesin-8X is located on the spindle during mitotic and meiotic divisions at various stages of parasite life cycle. Deletion of this gene results in impaired endomitotic replication in oocyst development and sporogony in mosquitoes, thereby blocking transmission of the parasite to its vertebrate host.

## Results

### Phylogenetic analysis of kinesins in Apicomplexa identifies 15 families of which nine are present *Plasmodium*

Using a previously published dataset of kinesin protein sequences as a starting point [3], we conducted a bioinformatic analysis to produce a high-resolution phylogenetic distribution of all known apicomplexan kinesins. We detected nine kinesin genes in *P. berghei*, including the two Kinesin-8 genes (Fig 1, S1 Fig and S1 Table1). Kinesin-8X is evolutionary conserved across all apicomplexan genomes whereas Kinesin-8B is restricted to Haemosporidia and Cocccidia. Remarkably, we do not detect Kinesin-4 in the Laverenia species, *P. falciparum* and *P. reichenowi* and this was further verified by comparison of orthologous genes at the syntenic regions within the genomic context at chromosomal levels between *P. berghei* ANKA and *P. falciparum* 3D7 genomes.

**Fig 1.**
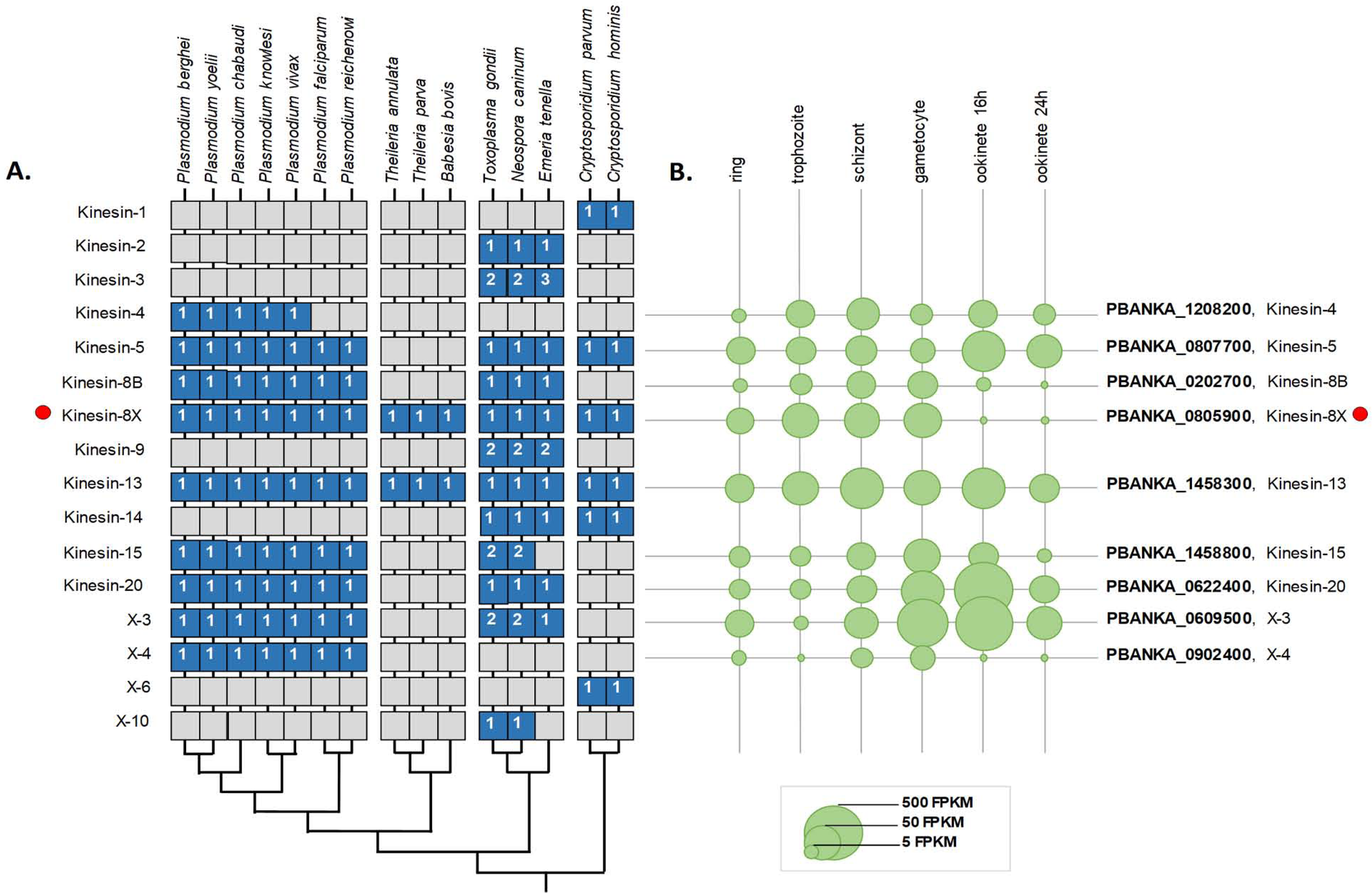
Phylogenetic analysis of apicomplexan kinesins. **(A)** Phylogenetic distribution of detected kinesin genes in alveolate genomes. Blue boxes denote the presence of genes, with the number of detected genes shown. **(B)** The expression levels of *P. berghei* genes in different developmental stages [39] are shown as circles. Note that *Plasmodium spp* contain two Kinesin-8 genes.

Similar to Kinesin-8X, Kinesin-13 genes are conserved in evolution across all Apicomplexa clades – suggesting essential roles of these kinesin genes. With the exception of coccidian Kinesin-9, Kinesin-15, and Kinesin-X3, multiple kinesins detected within a single genome all belong to the same orthogroup (http://orthomcl.org/orthomcl/), suggesting that they arose derived through gene duplication. Our analysis revealed an apparent scarcity of kinesins genes in piroplasm genomes (Fig 1, S1 Fig and S1 Table1), which may have resulted from highly divergent piroplasm kinesins not easily identified by comparative studies with existing bioinformatic protocols.

### Kinesin-8X motor domain has MT-based motility and depolymerization activities

We were interested to investigate the roles of Kinesin-8s in *Plasmodium* especially with respect to regulation of microtubule dynamics during cell division, focusing on Kinesin-8X. To understand the molecular properties of the proteins encoded by *P. berghei* (*Pb)* Kinesin-8X (PBANKA_0805900) and its orthologue in *P. falciparum* (*Pf*, PF3D7_0319400), we began by studying the biochemistry of their conserved motor domains. Both motor domains are located in the middle of the protein sequences (Fig 2A) and show 91% sequence identity. We expressed these motor domains as recombinant proteins – referred to below as *Pb*Kinesin-8X-MD and *Pf*Kinesin-8X-MD, respectively - and characterised their activities in standard kinesin assays. *Pb*Kinesin-8X-MD and *Pf*Kinesin-8X-MD exhibit MT-stimulated ATPase activity (Fig 2B). The V_max_ values are 1.5 and 5.7 ATP/s for *Pb*Kinesin-8X-MD and *Pf*Kinesin-8X-MD, respectively, and their K_1/2(MT)_ are 2.2 and 1.4 μM, respectively. These motors are therefore slow ATPases with weaker affinity for MTs compared to for example the human Kinesin-8 KIF18A motor domain [38].

**Fig 2.**
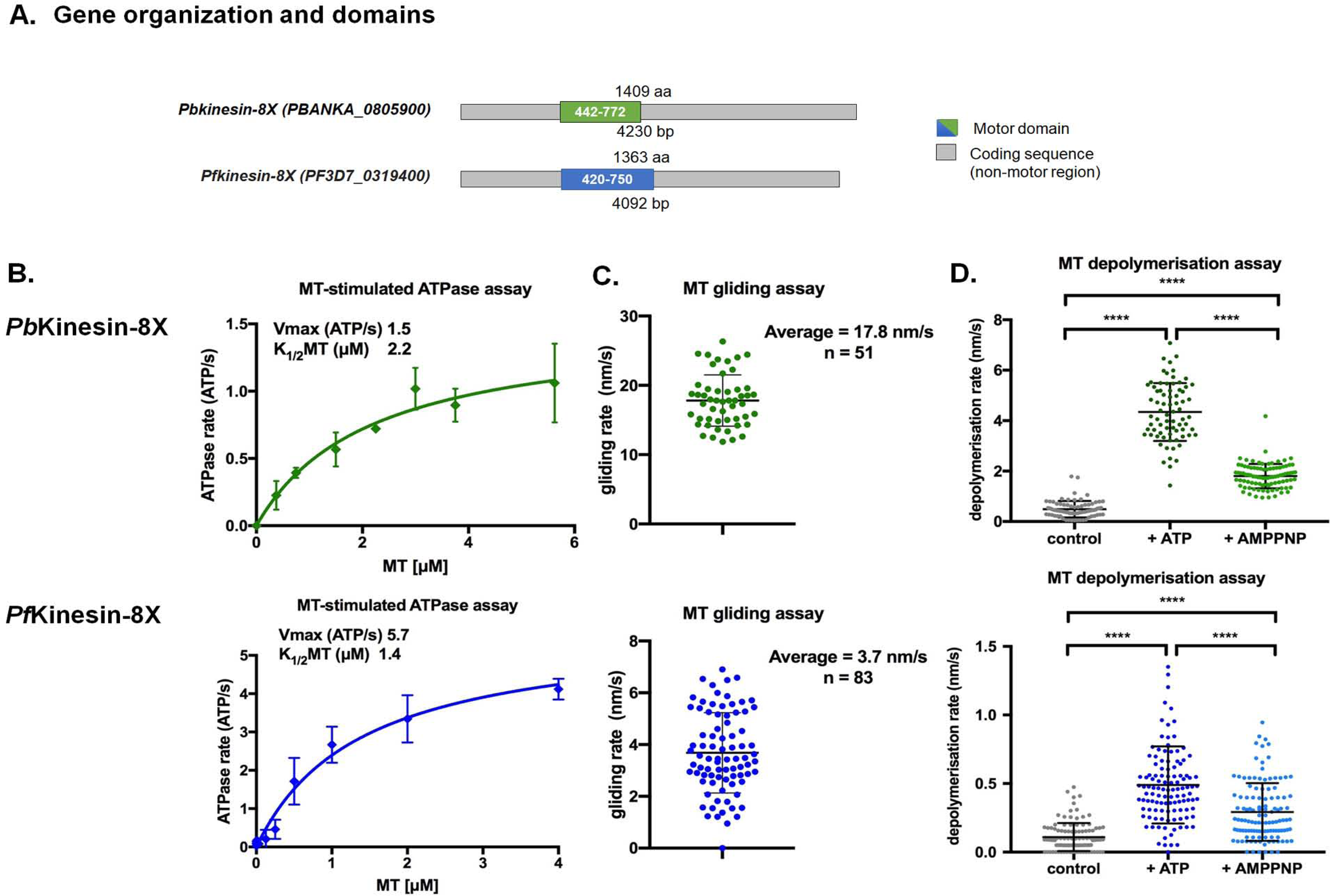
Kinesin-8X shows ATPase, gliding motor and depolymerization activities. **(A)** Schematic protein organisation of PBANKA_0805900 (PbKinesin-8X) and PF3D7_0319400 (PfKinesin-8X) showing their full-length sequence and central location of the motor domain, PbKinesin-8X (green) and PfKinesin-8X (blue). **(B-D)** Activities of Pb (top - green) and Pf (bottom – blue) Kinesin-8X motor domains in three kinesin assays. **(B)** Microtubule (MT) stimulated ATPase activity; data fitted to Michaelis-Menten equation with calculated V_max_ and K_m_ parameters. **(C)** Microtubule gliding activity measured by TIRF microscopy; the average motility (nm/s) and the number of microtubules measured are shown. The difference between PbKinesin-8X and PfKinesin-8X velocity is statistically significant (t-test P <0.0001). **(D)** Microtubule depolymerization measured using TIRF microscopy; depolymerization rate (nm/s) in the presence of ATP and AMPPNP is compared to a control in the absence of nucleotide. t-tests were performed in Prism to establish the significance of the nucleotide-dependent differences and differences between species. Significance values are displayed as asterisks, all p-values were <0.0001 (****) comparing control with the presence of AMPPNP or ATP and comparing activity in the presence of AMPPNP or ATP.

Both Kinesin-8X motor domains drive MT gliding (Fig 2C), with an average velocity of 17.8 ± 3.7 nm/s (n = 51) and 3.7 ± 1.7 nm/s, (n = 54) for *Pb*Kinesin-8X-MD and *Pf*kinesin-8X-MD, respectively. These velocities are less than that of an equivalent human Kinesin-8 KIF18A motor domain, with a gliding velocity of ~40 nm/s [38]. Both Kinesin-8X motor domains depolymerized paclitaxel-stabilised MTs, a characteristic of a Kinesin-8 subset from other species [17, 18] (Fig 2D). Significant MT depolymerization was observed in the presence of AMPPNP or ATP, compared to the no nucleotide control. *Pb*Kinesin-8X-MD (4.3 ± 1.1 nm/s +ATP, n = 74; 1.8 ± 0.5 nm/s +AMPPNP, n = 93) depolymerised MTs faster than *Pf*Kinesin-8X-MD (0.5 ± 0.3 nm/s +ATP, n = 116; 0.3 ± 0.2 nm/s +AMPPNP, n = 117). Similarly, for both proteins, depolymerization was faster in the presence of ATP than AMPPNP. In summary, we conclude that the *P. berghei* and *P. falciparum* Kinesin-8X motor domains are comparatively slow microtubule translocators that depolymerize microtubules *in vitro* and hence have the potential to regulate MT dynamics *in vivo*.

### Kinesin-8X is transcribed at most *P. berghei* developmental stages

To understand the context in which these activities could operate, we investigated expression and localisation of Kinesin-8X in *P. berghei* by first examining its transcript level using quantitative real time PCR (qRT-PCR) at different developmental stages. The transcription profile showed expression of *Pb*Kinesin-8X at most stages of parasite development with the highest RNA level in gametocytes, followed by blood stage schizonts and ookinetes (S2A Fig). These results are comparable to those obtained in previous RNA-seq analyses of *P. berghei* (Fig 1B) [39, 40].

### Spatio-temporal profile of Kinesin-8X revealed by live cell imaging of mitotic and meiotic stages in parasite development

To investigate the subcellular location of Kinesin-8X in *P. berghei*, we generated a transgenic parasite line by single crossover recombination at the 3’ end of the endogenous *kinesin-8X* locus to express a C-terminal GFP-tagged fusion protein (S2B Fig). PCR analysis of genomic DNA using locus-specific diagnostic primers indicated correct integration of the GFP tagging construct (S2C Fig). The presence of protein of the expected size (~188 kDa) in gametocyte lysate was confirmed by western blot analysis using GFP-specific antibody (S2D Fig). The Kinesin-8X-GFP parasites completed the full life cycle with no detectable phenotype resulting from the GFP tagging (data not shown).

The expression and localization of *Pb*Kinesin-8X was assessed by live cell imaging throughout the parasite life cycle. *Pb*Kinesin-8X was not detectable by microscopy in asexual blood stages (S2E Fig) but was observed in both male and female gametocytes, with a diffuse nuclear localization. Following activation of gametogenesis with xanthurenic acid and decreased temperature *in vitro* [28, 41]. *Pb*Kinesin-8X protein began to accumulate in male gametocytes at one end of the nucleus, presumably at the putative MTOC. Within one-minute of activation we observed the distribution of Kinesin-8X, like an arc across the nucleus, later forming two distinct foci that could be consistent with the formation of two MTOCs (Fig 3A). As DNA replication and endomitosis continued in male gametogenesis, six to eight distinct foci were seen to form 8 to 10 min after activation. These Kinesin-8X foci could be associated with MTOCs of the 8N nucleus that precedes exflagellation to produce eight male gametes (Fig 3A). There was no detectable expression of PbKinesin-8X in these male gametes. To examine further the location of PbKinesin-8X we investigated its co-localization with MTs (using α-tubulin as a marker) by indirect immunofluorescence assay (IFA) and fixed cells. PbKinesin-8X was localized on MTs during the early stages of male gametogenesis but in later stages it was distributed diffusely within the nucleus (Fig 3C). To improve visualisation, we used deconvolution microscopy and confirmed that Kinesin-8X is localized on mitotic spindles in early stages of male gametogenesis. (Fig 3D). In female gametocytes Kinesin-8X showed no major change in distribution and remained nuclear even 15 min post activation (Fig 3B). During this period the nucleus, and the pattern of Kinesin-8X, became more condensed and centrally located within the female gamete.

**Fig 3.**
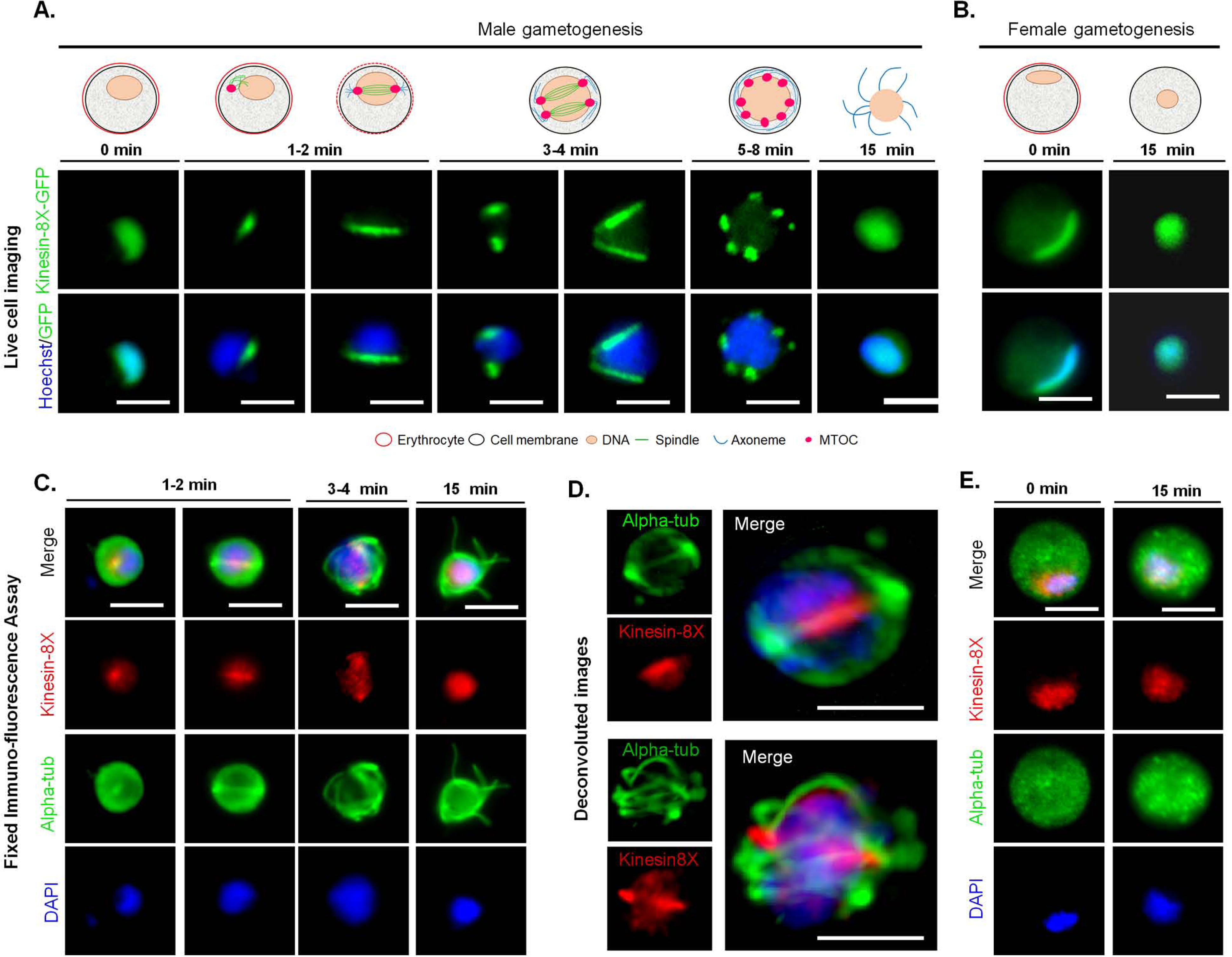
Dynamics of *Pb*Kinesin-8X show localization on spindle fibres during male gametogenesis. **(A)** Live imaging of *Pb*Kinesin-8X-GFP (Green) during male gametogenesis showing an initial location on putative microtubule organizing centre (MTOC) just after activation, and then on spindles and spindle poles in later stages. **(B)** Localization of *Pb*Kinesin-8X (red) during female gametogenesis before (0 min) and after activation (15 min). **(C)** Indirect immunofluorescence assays showing co-localization of *Pb*Kinesin-8X (red) and α-tubulin (green) during male gametogenesis. **(D)** Deconvoluted images of male gametocytes showing *Pb*Kinesin-8X (red) with α-tubulins (green). **(E)** Indirect immunofluorescence assays showing location of *Pb*Kinesin-8X (red) and α-tubulin (green) during female gametogenesis. Scale bar=5 μm.

Next, we examined the location and dynamics of Kinesin-8X during ookinete development from the zygote. The zygote, formed following fertilization, develops further into the motile ookinete over 24 h [42]. Two hours after fertilisation, Kinesin-8X began accumulating at one end of the nucleus, as determined by live cell imaging. Following the initial protrusion of apical membrane during stage I and II of ookinete development, Kinesin-8X-GFP was observed on spindles. In later stages (stage V) it accumulated at two distinct foci, probably at two spindle poles, and remained there in the mature stages of ookinete development (Fig 4A). Interestingly, some early mature ookinetes (stage V-VI) also displayed some Kinesin-8X located at the basal end of the cell, but it disappeared from this location in fully mature ookinetes (Fig 4B).

**Fig 4.**
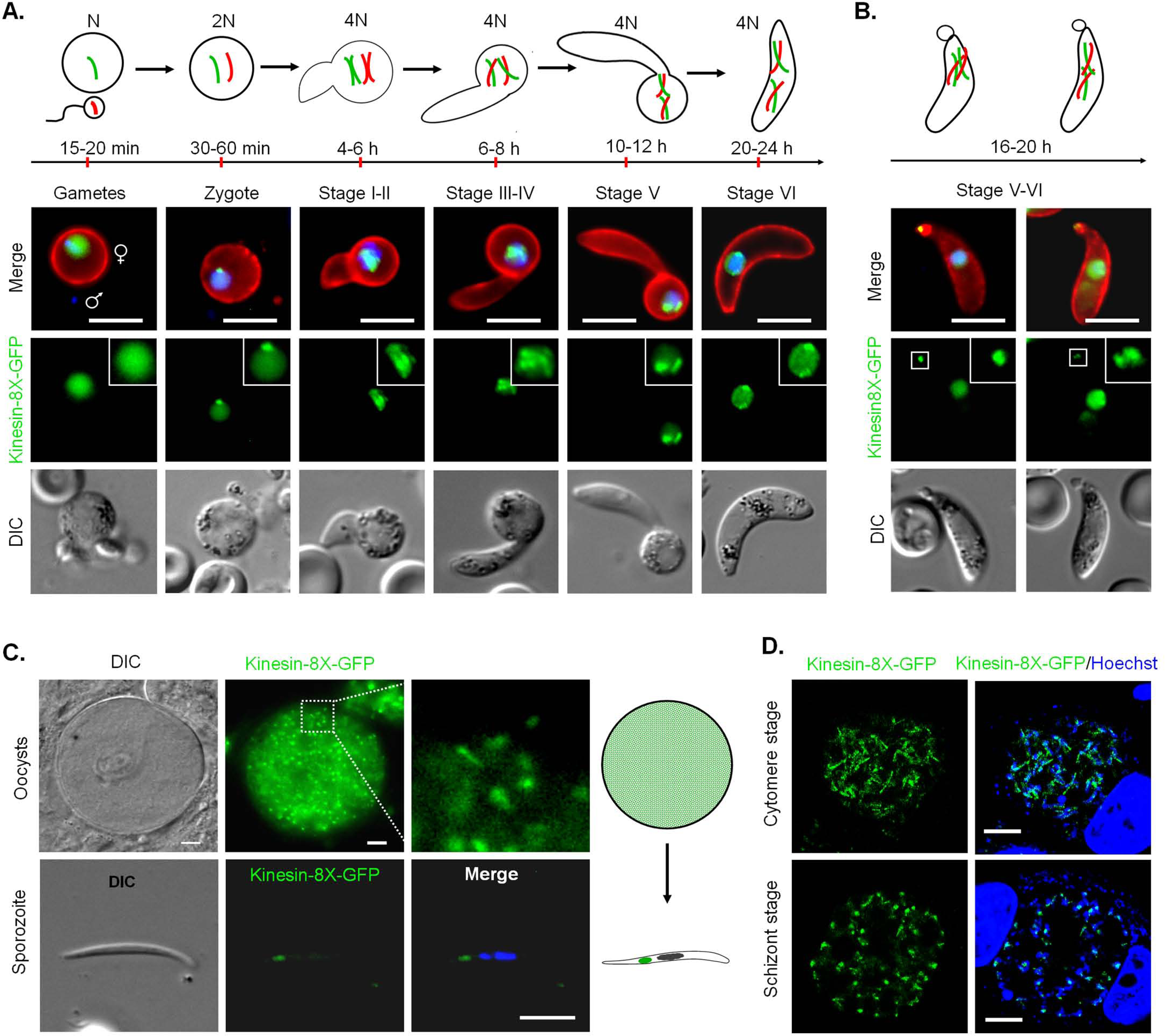
*Pb*Kinesin-8X localizes to a putative MTOC and spindle during ookinete development, sporogony and liver stage development. **(A)** Live cell imaging showing *Pb*Kinesin-8X-GFP location during ookinete development. A cy3-conjugated antibody, 13.1, which recognises the protein P28 on the surface of activated female gametes, zygotes and ookinetes was used to mark these stages (red). **(B)** Representative images showing *Pb*Kinesin-8X, located at basal end of early-mature ookinetes as well as in the nucleus. **(C)** *Pb*Kinesin-8X-GFP location in oocyst and sporozoite. **(D)** Location of *Pb*Kinesin-8X in liver stages. Scale bar=5 μm.

Ookinetes traverse the mosquito gut epithelium and develop into oocysts that produce sporozoites. During oocyst development at about 10 to 14-days post mosquito feeding, Kinesin-8X was observed as punctate dots located near putative MTOCs (Fig 4C). Many arc or bridge-like structures were also observed that may represent the movement of Kinesin-8X on spindles during this endomitosis in oocysts. *Pb*Kinesin-8X was also present in sporozoites, located as a focal point next to the nucleus (Fig 4C). We further analysed Kinesin-8X-GFP location by crossing with Ndc80-Cherry line in oocyst and sporozoites. Kinesin-8X was observed close to the nucleus and adjacent to Ndc80, but does not totally overlap with it in both oocyst and sporozoites.

The sporozoites were used to infect HeLa cells, to study the expression and location of *Pb*Kinesin-8X during the vertebrate pre-erythrocytic stage. The pattern of protein localization during liver stage development was similar to that in other mitotic stages, showing a spindle pattern in cytomere stages and an MTOC like location in schizont stages (Fig 4D).

### *Pb*Kinesin-8X is required for endomitotic division during oocyst development and for transmission of the parasite

To assess the importance and function of Kinesin-8X throughout the *Plasmodium* life cycle, the *P. berghei* gene was deleted using a double crossover homologous recombination strategy (S2F Fig). Diagnostic PCR was performed to confirm successful integration of the targeting construct at the *kinesin-8X* locus (S2G Fig). Analysis of these transgenic parasites by qPCR confirmed complete deletion of the *kinesin-8X* gene (S2H Fig). The successful deletion of the *kinesin-8X* gene indicates that it is not essential during the asexual blood stage of parasite development and this idea is supported by a recent study in which functional profiling of the *Plasmodium* genome was performed [43]. Phenotypic analysis was then carried out at other developmental stages of the two independent clones of *Δkinesin-8X* parasite, comparing it with the wild type (WT-GFP) parasite as a control. Both knockout clones showed same phenotype and data presented here is a combined data for both clones. There was no difference in exflagellation during male gametogenesis (Fig 5A), and zygote formation and ookinete development were also not affected (Fig 5B). To assess the effect on oocyst development, *Anopheles stephensi* mosquitoes were fed on mice infected with *kinesin-8X* parasites, and the number of GFP-positive oocysts on the mosquito gut was counted 7 days later. There was no significant difference in the number of *Δkinesin-8X* oocysts compared to WT controls at 7 dpi (days post infection), however a significant reduction was observed at 10 dpi, which became even more evident at 14 (Fig. 5C). By day 21 post infection the number of GFP-positive *Δkinesin-8X* oocysts had decreased further to only 8-10% of the WT-GFP parasite number (Fig 5C). The *Δkinesin-8X* oocysts were approximately half the size of WT-GFP oocysts at day 10 and 14, and even smaller at day 21 post-infection (Fig 5D). Most of the *Δkinesin-8X* parasites oocysts were dead on day 21 showing less GFP expression and disintegrated nuclei (Fig 5E) and no viable sporozoites were observed in oocysts (Fig 5F). We also verified for sporozoites in salivary glands and found no *Δkinesin-8X* parasites (Fig 5G). As there were no viable sporozoites in salivary glands, mosquitoes infected with *Δkinesin-8X* parasites did not transmit the parasite to susceptible mice. In contrast, mosquitoes infected at the same time with WT-GFP parasites were able to transmit the disease and blood stage infection was observed in naïve mice 4 days later (Fig 5H).

**Fig 5.**
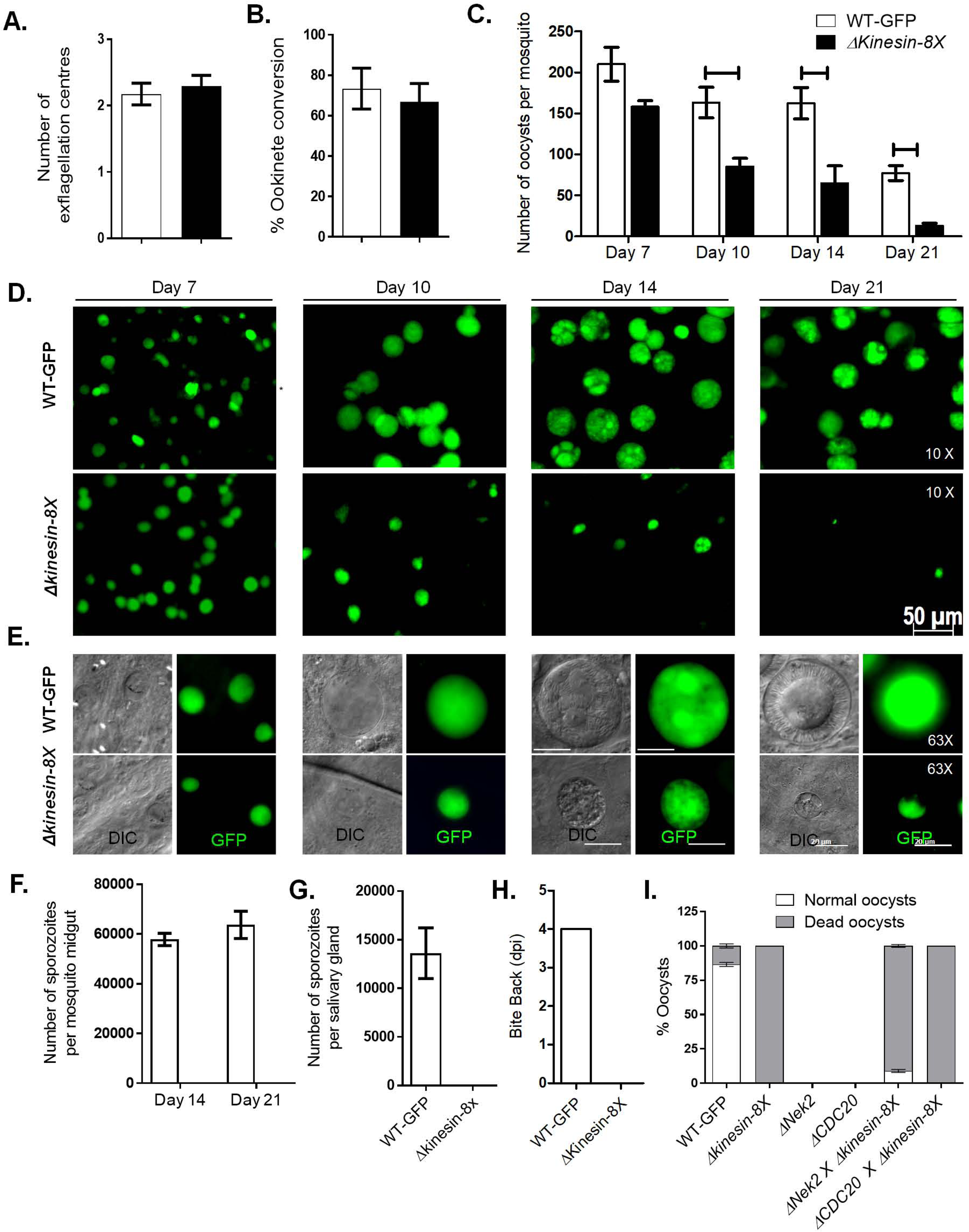
*Pb*Kinesin-8X is essential for oocyst development and sporogony. **(A)** Male gametogenesis (exflagellation) of *Δkinesin-8X* line (black bar) compared with WT-GFP line (white bar) measured as the number of exflagellation centres per field. Mean ± SD. n=5 independent experiments. **(B)** Ookinete conversion as a percentage for *Δkinesin-8X* (black bar) and WT-GFP (white bar) parasites. Ookinetes were identified using 13.1 antibody as a surface marker and defined as those cells that differentiated successfully into elongated ‘banana shaped’ ookinetes. Mean ± SD. n=5 independent experiments. **(C)** Total number of GFP-positive oocysts per infected mosquito in *Δkinesin-8X* (black bar) compared to WT-GFP (white bar) parasites at 7, 10,14 and 21-day post infection (dpi). Mean ± SD. n= 3 independent experiments (>15 mosquitoes for each) *p≤0.05, **p≤0.01. **(D)** Mid guts at 10x magnification showing oocysts of *Δkinesin-8X* and WT-GFP lines at 7, 10, 14 and 21 dpi. Scale bar=50 μM. * *p* ≤ 0.05 and ** *p* ≤ 0.01 **(E)** Mid guts at 63x magnification showing oocysts of *Δkinesin-8X* and WT-GFP lines at 7, 10, 14 and 21 dpi. Scale bar= 20 μM. **(F)** Total number of sporozoites in oocysts of *Δkinesin-8X* (black bar) and WT-GFP (white bar) parasites at 14 and 21 dpi. Mean ± SD. n= 3 independent experiments. **(G)** Total number of sporozoites in salivary glands of *Δkinesin-8X* (black bar) and WT-GFP (white bar) parasites. Bar diagram shows mean ± SD. n= 3 independent experiments **(H)** Bite back experiments showing no transmission of *Δkinesin-8X* parasites (black bar) where WT-GFP parasites (white bar) show successful transmission from mosquito to mice. Mean ± SD. n= 3 independent experiments **(I)** Rescue experiment showing male allele of *Δkinesin-8X* is affected.

### Kinesin-8X function during sporogony is contributed by the male gamete

Since the Kinesin-8X is expressed in both male and female gametocytes and the parasite development is affected after fertilization, we investigated whether the defect is due to the inheritance of the male- or female gamete. We performed genetic crosses between *Δkinesin-8X* parasites and *P. berghei* mutants deficient in production of either male *(Δcdc20)* or female *(Δnek2)* gametocytes. Crosses between *Δkinesin-8X and Δnek2* mutants produced a few normal sized oocysts that were able to sporulate, showing partial rescue of the *Δkinesin-8X* phenotype (Fig 5I). On the other hand, crosses between *Δkinesin-8X and Δcdc20* showed no rescue of the *Δkinesin-8X* phenotype. These results reveal that a male copy of a functional kinesin-8X gene is required for oocyst development.

### Ultrastructure of Δ*kinesin-8X* parasites show defects in oocyst growth and sporozoite budding

To define further the defect in oocyst growth, midguts at 14 days post infection in both Δ*kinesin-8X* and WT-GFP parasite-infected mosquito were examined by transmission electron microscopy. In the WT-GFP parasites, numerous oocysts were observed at various stages of sporozoite development. Large numbers of sporozoites were observed budding from the cytoplasmic masses in WT-GFP parasites (Fig 6a, c). In contrast it was extremely difficult to identify any oocysts in Δ*kinesin-8X*. A detailed search identified a few collapsed oocysts with degenerate cytoplasmic organelles in Δ*kinesin-8X* parasites (Fig 6b, d, e).

**Fig 6.**
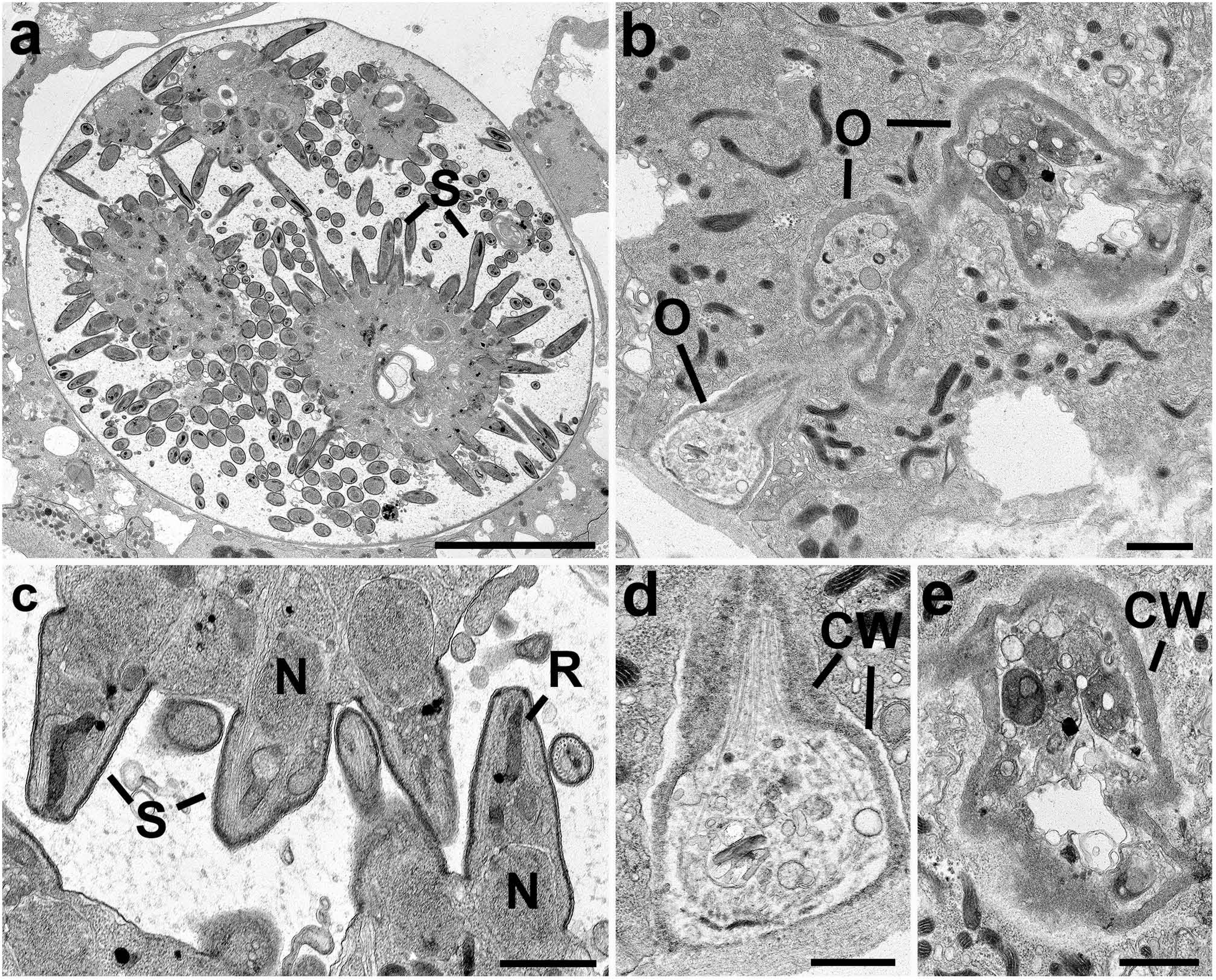
Ultrastructure analysis of oocyst development in Δkinesin-8X parasites. Electron micrographs of WT (a, c) and mutant (b, c, d) oocysts located in the mid gut of the mosquito at 14 days post-infection. Bar is 10 μm in a and 1μm in other micrographs. (a) Low power image through a mid-stage oocyst showing early stages in sporozoite (S) formation. (b) Low power showing the collapsed appearance of three oocysts (o). (c) Detail of an oocysts showing budding sporozoites (S) containing nucleus(N) and developing rhopty (R). (d, e) Enlargements of oocysts in image b showing the collapsed oocyst wall (CW) surrounding cytoplasm with degenerate organelles.

### Transcriptome analysis of Δ*kinesin-8X* parasites reveals modulated expression of genes involved in motor activity, and many other functions

Because oocyst development was affected in *Δkinesin-8X* parasites, we then analysed the expression of mRNA in *Δkinesin-8X* and WT-GFP parasites. Global transcription was investigated by RNA-seq analysis of WT-GFP and *Δkinesin-8X* gametocytes immediately before activation (0 min) and after exflagellation (30 min after activation). The genome-wide read coverages for the four pairs of biological replicates (WT, time 0; WT, time 30 min; *Δkinesin-8X*, time 0; and *Δkinesin-8X*, time 30 min) exhibited Spearman correlation coefficient of 0.97, 0.98, 0.95 and 0.95; respectively, validating the reproducibility of the experiment. The deletion of *kinesin-8X* in the *Δkinesin8X* strain was also confirmed in the RNA-seq analysis as no significant reads mapped to the region of this gene (Fig 7A). Expression of some other genes was either significantly upregulated or downregulated in the knockout strain. In total, 482 genes were upregulated, and 277 genes were downregulated in comparison to the WT-GFP control (Fig 7B, S3 Table). Gene ontology enrichment analysis showed that the expression of genes involved in MT-based processes - including motor activity - together with several other functions like cell division and chromosome organization were specifically upregulated, indicating a possible mechanism of compensation during cell division (Fig 7C). Three of the top 20 and six of top 50 most highly upregulated genes in the *ΔKinesin8X* strain encode putative dynein heavy chain proteins, further supporting a strong increase in expression of microtubule-related motor proteins and suggesting a specific association between dynein and kinesin (S3 Table).

**Fig 7.**
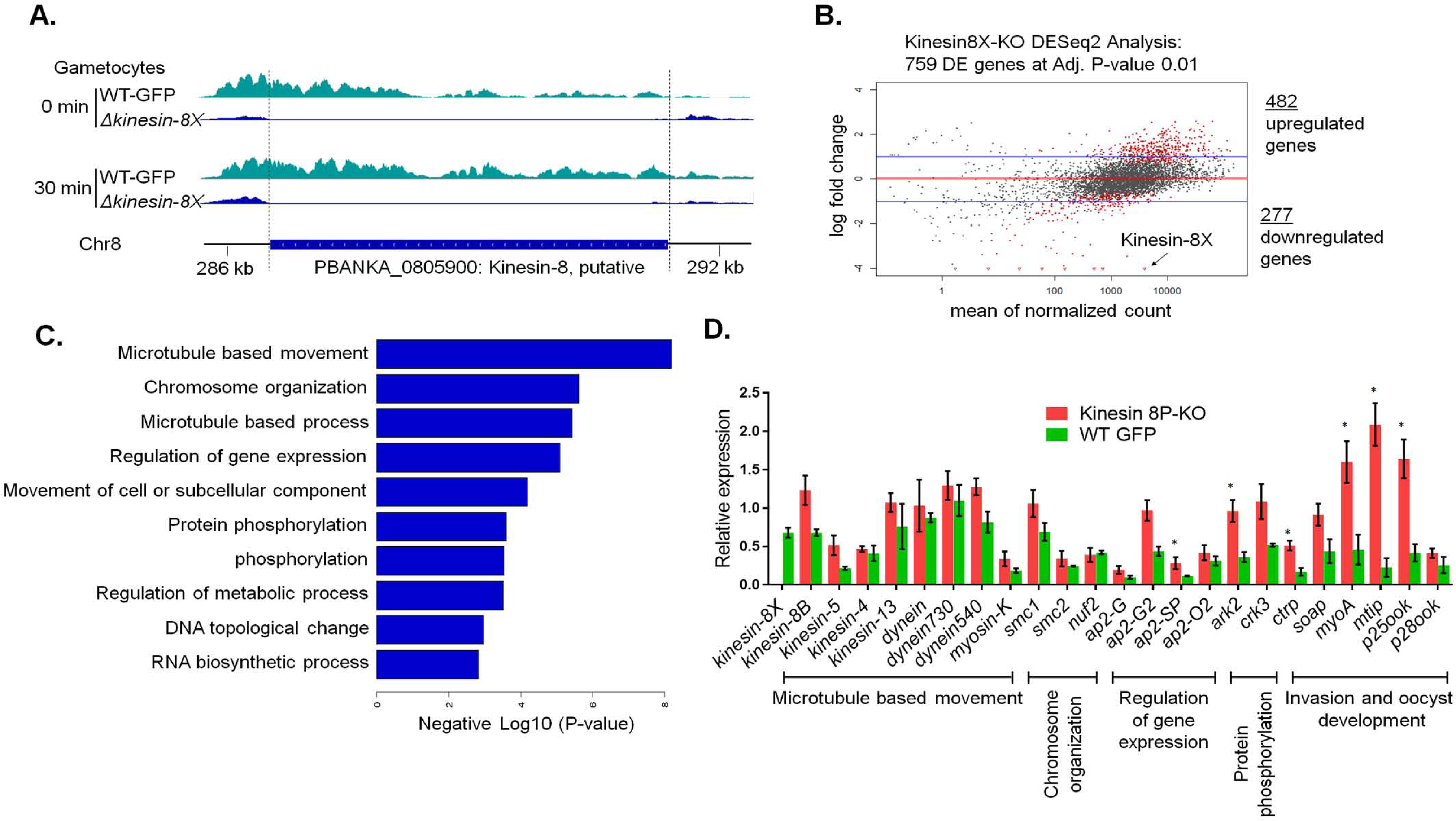
Global transcript analysis of *Δkinesin-8X* parasites by RNA-seq. **(A)** RNA sequence analysis showing no transcript in *Δkinesin-8X* parasites. **(B)** Upregulated and downregulated genes in *Δkinesin-8X* parasites compared to WT-GFP parasites. **(C)** Gene ontology enrichment analysis showing most affected genes involved in various biological processes. **(D)** Validation of relevant and selected genes from the RNA-seq data by qRT-PCR. Mean ± SD. n= 3 independent experiments. *p ≤ 5

Transcript levels detected by RNA-seq experiment were further validated by qRT-PCR for genes involved in motor activity, ookinete invasion and oocyst development (Fig 6D). Data showed good correlation with the RNA-seq results (Fig 7D) further validating the notion of a compensatory mechanism in this particular pathway.

## Discussion

The importance of Kinesin-8 molecular motors in mitosis in many eukaryotes is well-established. These motors move on spindle MTs towards their plus end and regulate spindle dynamics, contributing to spindle length and positioning, and thus to chromosome alignment during metaphase [44–46]. Although the exact molecular basis of these functions remains controversial, Kinesin-8s modulate MT dynamics and may either stabilise, destabilise, slow growth, or actively depolymerize MTs in spindle function [18, 19, 47]. Phylogenetic analysis showed that while there are 9 kinesin genes in *Plasmodium*, a number of kinesin classes, typically involved in long-distance intra-cellular transport in metazoa, such as kinesin-1 and kinesin-3, are absent [3, 37]. The phylogenetic analysis of Apicomplexa in this study identifies 15 Kinesin families, from which 9 Kinesins are encoded by *Plasmodium* genome. This analysis further supports the previous finding that the genome of *Plasmodium* encodes two Kinesin-8 proteins that have been classified as Kinesin-8B and Kinesin-8X subtypes. Only Kinesin-8X and Kinesin-13 are found throughout all investigated Apicomplexa. This observation could either reflect a high level of sequence diversity among Kinesin proteins,or underline the plasticity of Kinesin functions during the evolution of Apicomplexa genomes.

Protein sequence analysis of PbKinesin-8X, and its orthologue PfKinesin-8X, identified the conserved kinesin motor domain in the middle of the protein, in contrast to its N-terminal location in Kinesin-8s of most other organisms [44]. Nevertheless, expressed recombinant motor domains had MT-dependent gliding activity, like that of other Kinesin-8 motor domains, albeit slower than human Kinesin-8 Kif18A [38]. Such gliding activity of motors in an N-terminal location is usually indicative that a dimeric full-length kinesin can take multiple steps along MTs. Whether or not the central location of the motor domain modifies the behaviour or function of the full-length proteins will be the topic of future studies. The *Plasmodium* Kinesin-8X motor domains also showed MT depolymerisation activity, a characteristic shared with Kinesin-8 from yeast and humans [10, 17, 18, 38]. It is also interesting to note that while the MT-stimulated ATPase activity of PbKinesin-8X is slower than that of PfKinesin-8X, its ATP-dependent gliding activity and ATP-dependent rate of MT depolymerisation are faster therefore, understanding the origin of the apparent inefficiency of our PfKinesin-8X recombinant protein compared to PbKinesin-8X will also require further work. PbKinesin-8X and PfKinesin-8X-catalysed MT depolymerisation is faster in the presence of ATP than in the presence of AMPPNP; this differentiates them from human Kinesin-8 [38] and could be indicative of differences in depolymerisation mechanism between Kinesin-8s in these species. However, the overall biochemical properties of the Plasmodium Kinesin-8X motor domains are consistent with the idea that these proteins combine activities of stepping and MT dynamics control, to ensure accurate cell division in these parasites.

Microscopy of PbKinesin-8X suggests it is located at the putative MTOC and on the spindle fibres in most of the proliferative stages found within the mosquito vector including male gametogenesis, ookinete differentiation and sporogony. Male gametogenesis is a very rapid process with three rounds of DNA replication within 8 to 10 min of activation, associated with three endomitotic divisions and the assembly of eight flagella, leading to exflagellation, the release of eight male gametes [27, 29]. The dynamic localization of Kinesin-8X mirrors MT dynamics (cytoskeleton arrangements) during the three rounds of replication. The colocation of PbKinesin-8X and α-tubulin was shown using specific antibodies and confirms Kinesin-8X localization on spindles and MTs during chromosome replication in male gametogenesis. A similar location and role in MT dynamics has been shown in fission yeast [48], budding yeast [49], Drosophila [50], and human [10] where Kinesin-8 proteins are co-localized with MTs during cell division. The absence of kinesin-8X indicates that it is not associated with MTs in the motile gamete which suggests the spindle /MTOC rapidly depolymerise after exflagellation.

The discrete foci of Kinesin-8X during zygote development and differentiation suggest it is associated with the putative MTOC at this stage, when the genome is replicated from 2N to 4N. This is the stage of parasite development when meiosis and genetic recombination occur [51], and after the succeeding oocyst formation, finally giving rise to haploid sporozoites. The spatio-temporal profile of Kinesin-8X during the zygote to ookinete transition (stage 1-VI) suggest a role in chromosome replication during meiosis. The additional location of PbKinesin-8X at the basal end of early mature ookinetes (stage V-VI) and its disappearance in fully mature ookinetes is also consistent with a role in movement of the nucleus from the cell body to the ookinete. A somewhat similar role of Kinesin-8 is in positioning the nucleus at the centre of the cell in fission yeast [20, 22]. The mature ookinete crosses the mosquito gut epithelium and on the basal side of the gut wall it rounds up and develops into an oocyst. Endomitosis in the oocyst consists of many rounds of nuclear division to produce thousands of sporozoites [31, 52]. The distinct Kinesin-8X-GFP foci in early stage oocysts and localization to the MTs suggest that these foci are MTOCs and bridges are the spindles. This suggestion is supported by earlier EM studies of oocysts, in which hemi-spindles and MTOCs were found in the developing oocyst stage [31, 52, 53].

Expression of Kinesin-8X in the pre-erythrocytic liver stages suggests that it is also involved in this endomitotic stage of parasite replication although surprisingly, it is apparently not present in blood stage schizogony; it was not detected by either live cell imaging or fixed cell immunofluorescence. It is fascinating that the asexual blood stages are negative suggesting there may be alternate motors at this stage of the life cycle. However, all together our localization data suggests an important role of Kinesin-8X in the regulation of MT dynamics during DNA replication and chromosome segregation in endomitosis and meiosis during asexual and sexual stages.

The deletion of *pbkinesin-8X* had no phenotype during asexual multiplication in either the erythrocytic phase, in male gametogenesis, or during ookinete development suggesting that during these stages of parasite development this motor is not essential. It is also most likely that other kinesins can compensate at these particular stages for the loss of PbKinesin-8X, perhaps most importantly this could involve Kinesin-8B, the second motor protein of the same family in *Plasmodium* [3] (Zeeshan et al unpublished observation). The deletion of Kinesin-8 in budding yeast and *Drosophila* results in uncontrolled spindle growth, with long metaphase spindles and delayed spindle shortening in anaphase [15]. Similar consequences have been observed in *Xenopus* upon depletion of Kinesin-13 [54], suggesting that Kinesin-8 and Kinesin-13 have similar depolymerization activities that can compensate for the loss of each other. The *Plasmodium* genome encodes a Kinesin-13 that has depolymerization activity in vitro [55], and the upregulation of Kinesin-13 and Kinesin-8B in the Δ*kinesin-8X* parasite might suggest that the loss of this gene is compensated by overexpression of either Kinesin-8B or Kinesin-13 or both at most stages of the life cycle.

Despite a lack of phenotype at other stages, oocyst formation and maturation were impaired in *Δkinesin-8X* parasites, producing fewer oocysts of smaller size when compared to WT-GFP parasites, in the mosquito gut. Further in-depth analysis of oocysts in midgut of *Δkinesin-8X* parasites by electron microscopy revealed that division of nuclear lobes with oocyst growth is drastically affected and further budding of sporozoites from these lobes does not occur. Plasmodium oocysts contain a highly lobed syncytial nucleus that divides at the time of sporozoite budding into a number of lobes, which undergo subsequent mitotic division resembling endomitosis that was observed in WT-GFP parasites. This suggests that Kinesin-8X is involved in the correct onward differentiation of invasive ookinetes and the endomitotic process during sporogony. A similar phenotype was observed in our recent study on *Plasmodium* specific P type cyclin PbCYC3 during sporogony [56], in which oocyst size and sporozoite formation are affected. Similar results were observed with other gene deletion mutants including MISFIT [57], PK7 [58], DMC1 [59] and PPM5 [60]. The partial compensation of the Δ*kinesin-8X* phenotype with a female line mutant (*Δnek2)* indicates that the male lineage is affected in the parasite. This resembles what was observed for Δ*misfit* and *Δppm5*, both of which have an absolute requirement for a functional gene from the male line [57, 60]. In a genome-wide transcript analysis of the *Δppm5* phosphatase-deficient line, kinesin-8X was upregulated in activated gametocytes [60], suggesting that PPM5 may regulate the activity of Kinesin-8X by dephosphorylation. The morphology and DNA content of Δ*kinesin-8X* ookinetes is not different from that of WT-GFP parasites, as was also observed for *Δppm5* parasites, and in contrast to *Δmisfit* parasites in which the ookinete DNA content was less than in WT-GFP [57, 60]. One reason for fewer Δ*kinesin-8X* oocysts may be a defect in ookinete invasion of the mosquito gut wall, as shown recently for *Δppl4*, a deletion of the *Plasmodium* perforin-like protein-4 gene [61]. Interestingly, the expression of PPL4 was downregulated up to 50% in Δ*kinesin-8X* parasites. These findings, in conjunction with the data on the expression and localization of Kinesin-8X, suggest that this Kinesin has additional roles during oocyst development; most importantly, during the ookinete to oocyst transition.

Global transcript and qPCR analyses of the *Δkinesin-8X* parasites showed significant differential gene expression between knockout and WT strains. The genes that were mostly affected in Δ*kinesin-8X* parasites were involved in MT-based movement. An in-depth analysis of these genes showed upregulation of several kinesins, including *kinesin-8B, kinesin-13* and *kinesin-5*, suggesting that their activity might compensate for the loss of *kinesin-8X*. In addition, a number of genes encoding dynein heavy chains were among the most highly upregulated and one dynein light chain was downregulated, suggesting that kinesin and dynein may exhibit functional redundancy in which an increase in dynein expression can recover a specific function lost in the absence of Kinesin-8X [62]. Several other genes involved in transcriptional regulation such as AP2 transcription factors (e.g. AP2-O2, AP2G2, AP2SP) or gene involved in invasion or oocyst development were significantly upregulated, however it is important to highlight that modulation of these genes at the expression level was not able to recover the critical role of *kinesin-8X* during ookinete and oocyst development.

In conclusion, our work reveals that Kinesin-8X is associated with both mitotic and meiotic spindles during atypical cell division in all replicative stages of the life cycle with the notable exception of the asexual blood stages. Most importantly we validated the essential role of Kinesin-8X during endomitosis in oocyst development, indicating that specific inhibition of Kinesin-8X could be developed in novel transmission blocking strategies against these devastating malaria parasites.

## Methods

### Phylogenetic analysis of Apicomplexan Kinesins

The Hidden Markov Model (HMM) model for the kinesin motor domain (PF00225) was downloaded from Pfam (http://pfam.xfam.org/)[63] and searched against a range of apicomplexan genome-encoded protein data sets (Supp. Table 1). As in previous study [3], a preliminary threshold was set to 1e-25. To search for additional apicomplexan kinesins with divergent domains, we employed an additional search strategy using previously published kinesin sequences retrieved from *P. falciparum*, *T. annulata*, *T. gondii*, *C. parva*, and *T. thermophila* [3]. Reciprocal best BLASTP hits for these kinesins were obtained from an extended set of alveolate protein sets (Supp. Table 1), and for each Kinesins the corresponding sequences were collected. Sequences were aligned with mafft [64] and trimmed using trimAl [65]. Lineage-specific HMMs were then built using HMMer (http://hmmer.org/) and used to search the apicomplexan protein sets. To remove false-positives, a simple neighbour-joining tree was constructed (clustalw) from all proteins with detected domains, and all pair-wise genetic distances were calculated. For each protein, the average genetic distance to all other candidates for the same kinesin was compared to the average distance calculated individually for all other types of kinesin proteins. Only if the average distance to the same type of kinesin (for example, the average distance between a Kinesin-1 candidate and all other Kinesin-1 proteins) was lower than for all other kinesin types, were candidates retained. We refer to these two approaches as ‘direct HMM’ and ‘lineage-specific HMM’, respectively.

The direct HMM approach detected a group of Plasmodium and coccidian proteins with kinesin motor domains that showed no clear association to previously reported kinesins [3]. These proteins are not included in the presented sets but are listed in Supp. Table 2 as ‘Kinesin-like’. A maximum likelihood tree of the detected Kinesin proteins is shown in S1 Fig.

A total of 124 Kinesins were detected from the direct HMM approach, and 112 of these (90%) were also identified in through the lineage-specific HMM approach. The only additional proteins that were detected by the lineage-specific approach was a group of orthologous zinc-finger proteins (PBANKA_1351200, PY17X_1356300, PCHAS_1355800, PKNH_1263900, PVX_082800, PF3D7_1337400, PRCDC_1336400, TA20465, TP01_0517, BBOV_III002540, TGME49_269940, NCLIV_036650) that showed sequence similarity to X5 Kinesins from the Ciliate, *T. thermophila*. This group was deemed not to represent *bona fide* kinesins, as none of these proteins contained a Kinesin motor domain and sequences were divergent from all other kinesins.

### *P. berghei* Kinesin-8X recombinant protein expression and purification

DNA sequences-to express *P. berghei kinesin 8X* motor domain (residues 432-784, termed PbKinesin8X-MD) and an equivalent C-terminal SNAPf–tagged construct (Pbkinesin-8X-MD-SNAP) were cloned into a pNIC28-Bsa4 vector (Structural Genomics Consortium) including a TEV-cleavable N-terminal His_6_ tag using LIC cloning. The sequence was verified and the plasmid was transformed into *E. coli* BL21*(DE3) for protein expression.

Bacteria were grown at 37 °C until an OD_600_ of 0.7 and then switched to 20 °C for 30 min before addition of IPTG to 0.5 mM to induce protein expression. After 12 hours, cells were harvested and resuspended in lysis buffer (20 mM Tris-HCl pH 7.5, 500 mM NaCl, 5 mM MgCl_2_ and 1 mM ATP with EDTA-free protease inhibitor (Roche)). The cell suspension was sonicated for 45 min and then centrifuged at 48,384 g, 4 °C for 1 hr. The resulting supernatant was incubated with Ni-NTA agarose resin with mixing at 4 °C for 1 hr followed by washing with lysis buffer to reduce non-specific binding. The His_6_-PbKinesin-8X-MD constructs were eluted with lysis buffer containing 100mM Imidazole and incubated with TEV protease for 12 hrs to remove the tag. The protein was then exchanged by dialysis to low-salt ion exchange buffer (20 mM Tris-HCl pH 7.5, 100 mM NaCl, 5 mM MgCl_2_) and Ni-NTA resin was used to remove the His_6_-TEV protease from, the flow-through that contained PbKinesin-8X-MD without the His_6_-tag. Further purification was performed using a 1 ml HiTrap Q HP anion exchange column to remove any residual bacterial protein contaminants from the PbKinesin-8X-MD that did not bind. The Q column flow-through was concentrated, aliquoted and snap-frozen until further use.

### *P. falciparum* Kinesin-8X recombinant protein expression and purification

DNA sequences to express *P. falciparum kinesin-8X* motor domain (residues 420-762, termed PfKinesin-8X-MD) and an equivalent C-terminal SNAPf–tagged construct (PfKinesin-8X-MD-SNAP) were cloned into a pNIC-CTHF vector (Structural Genomics Consortium) that includes a TEV-cleavable C-terminal His_6_-FLAG tag using Gibson cloning. The DNA sequence was verified and the plasmid transformed into BL21*(DE3) *E. coli* cells for protein expression.

Bacteria were grown at 37 °C until the OD_600_ was around 0.6 to 0.8, then the culture was cooled to 18 °C before addition of IPTG to 0.1 mM to induce protein expression. After 12 hours, cells were harvested and resuspended in lysis buffer (50 mM Tris-HCl pH 7.0, 400 mM NaCl, 2 mM MgCl_2_, 1 mM ATP, 2 mM β-mercaptoethanol, 15 μg/ml DNase I (Sigma) with EDTA-free protease inhibitor (Roche). Cells were lysed using an Avesti Emulsiflex C3 high-pressure homogeniser, passaging the lysate three times. The lysate was centrifuged at 48,384 g, 4°C for 1 hr and the resulting supernatant was incubated with Ni-NTA agarose resin with mixing at 4 °C for 30 min followed by washing with low-imidazole containing buffer (50 mM Tris-HCl pH 7.0, 400 mM NaCl, 2 mM MgCl_2_, 1 mM ATP, 2 mM β-mercaptoethanol, 10 mM imidazole) to reduce non-specific binding. The His_6_-PfKinesin-8X-MD proteins were eluted with high-imidazole containing buffer (50 mM Tris-HCl pH 7.0, 400 mM NaCl, 2 mM MgCl_2_, 1 mM ATP, 2 mM β-mercaptoethanol, 250 mM Imidazole pH 7.0). Eluted fractions were dialysed for 12 hr at 4 °C against low-salt buffer (50 mM Tris-HCl pH 7.0, 40 mM NaCl, 2 mM MgCl_2_, 1 mM ATP, 2 mM β-mercaptoethanol) together with TEV protease to remove the C-terminal His_6_-FLAG tag. Dialysed protein was loaded onto a 1 ml HiTrap SP HP cation exchange chromatography column and eluted with gradient to a high-salt buffer (50 mM Tris-HCl pH 7.0, 1 M NaCl, 2 mM MgCl_2_, 1 mM ATP, 2 mM β-mercaptoethanol) on an ÅKTA system (GE Healthcare). PfKinesin-8X-MD containing fractions were pooled and loaded onto a Superdex 200 Increase 10/300 GL gel filtration column (GE Healthcare) equilibrated with gel filtration buffer (20 mM PIPES pH 6.8, 80 mM KCl, 2 mM MgCl_2_, 1 mM ATP, 2 mM β-mercaptoethanol). Fractions containing monomeric protein were pooled and concentrated to around 50 μM using Amicon Ultra-0.5 ml Centrifugal Filters (Millipore), and then aliquoted and snap-frozen until further use.

### MT polymerization

For all assays, porcine brain tubulin was purchased as a lyophilised powder (Cytoskeleton, Inc.) either unlabelled, X-rhodamine-labelled or biotinylated. The protein was solubilized in BRB80 buffer (80 mM PIPES-KOH pH 6.8, 1 mM EGTA, 1 mM MgCl_2_) to approximately 10 mg/ml (tubulin dimer concentration). Reconstituted tubulin was polymerised at 5 mg/ml final concentration in the presence of 5 mM GTP at 37 °C for 1 hr. After this a final concentration of 1 mM paclitaxel (Calbiochem) dissolved in DMSO was added and the microtubules incubated at 37 °C for another 1 hr.

#### ATPase assay

Unlabelled tubulin was polymerized as above in the presence of 5mM GTP and stabilized with paclitaxel. Free tubulin remaining after polymerization was removed by pelleting the microtubules by centrifugation at 392,000 g, removing the supernatant and resuspending the MT pellet in BRB80 buffer. Protein concentration was determined post-centrifugation by a Bradford assay.

#### Depolymerization assay

Microtubules containing 10% X-rhodamine-labelled and 10 % biotin-labelled tubulin (Cytoskeleton) were polymerized with GTP, paclitaxel-stabilised as above and left at room temperature for 48 hrs before use in a TIRF assay. **Gliding assay**: MTs containing 10% X-rhodamine-labelled tubulin were polymerized with GTP, paclitaxel-stabilised as above and left for 48 hrs at room temperature before use in a TIRF assay.

### ATPase activity

MT-stimulated kinesin ATPase activity was measured using a standard enzyme-coupled assay [66]. The assay was performed using 250 nM kinesin motor domain titrated with paclitaxel-stabilised MTs (0-6 μM) in 100 μl ATPase reaction buffer containing an ATP regeneration system (80 mM PIPES pH 6.8, 50 mM NaCl, 5 mM MgCl_2_, 1 mM EGTA, 5 mM phosphoenolpyruvate (PEP), 280 μM NADH, 12 U pyruvate kinase and 16.8 U lactate dehydrogenase). The ATP regeneration by pyruvate kinase is coupled to NADH depletion by lactate dehydrogenase in the conversion of PEP to lactate. NADH depletion was monitored by the decrease in absorbance at 340 nm in a SpectraMax Plus-384 plate reader every 30 s over 10 min at 37 °C (PbKinesin-8X-MD) and every 10 s for 1 hr at 26 °C (PfKinesin-8X-MD) operated by SoftMax Pro 5 software. The rates were corrected for the kinesin-only basal activity (PfKinesin8X-MD = 0.9 ATP/s; PbKinesin8X-MD = 0.7 ATP/s, plotted and used to calculate Km and Vmax using Prism software and fitting to the Michaelis-Menten equation.

### Microtubule depolymerization assay

Flow chambers for Total Internal Reflection Fluorescence (TIRF) microscopy were made between glass slides, biotin-PEG coverslips (MicroSurfaces Inc.), and double-sided tape. Chambers were sequentially incubated with: 1) blocking solution (0.75 % Pluronic F-127, 5 mg/ml casein) for 5 min, followed by two washes with assay buffer (80 mM PIPES, 5 mM MgCl_2_, 1 mM EGTA, 1 mM DTT and 20 μM paclitaxel); 2) 0.5 mg/ml neutravidin for 2 mins, followed by two washes with assay buffer (80 mM PIPES, 5 mM MgCl_2_, 1 mM EGTA, 1 mM DTT and 20 μM paclitaxel); 3) 1:100 dilution of X-rhodamine-labelled MT solution for 2 mins, followed by two washes with assay buffer supplemented with 1 mg/ml casein; 4) 2.5 μM unlabelled Kinesin8X-MD proteins in assay buffer supplemented with 5 mM nucleotide (as indicated) and an oxygen scavenging system (20 mM glucose, 300 μg/ml glucose oxidase, 60 μg/ml catalase). An Eclipse Ti-E inverted microscope was used with a CFI Apo TIRF 1.49 N.A. oil objective, Perfect Focus System, H-TIRF module, LU-N4 laser unit (Nikon) and a quad band filter set (Chroma). Movies were collected at room temperature under illumination at 561 nm for 30 min with a frame taken every 10 s with 100 ms exposure on a iXon DU888 Ultra EMCCD camera (Andor), using the NIS-Elements AR Software (Nikon). Where necessary, image drift was corrected using StackReg rigid body transformation. Depolymerisation rates were determined from kymographs using Fiji software. The assay was run in the presence of ATP, AMPPNP or apyrase as a no-nucleotide control. For each condition, data from two or more movies were analysed.

### Microtubule gliding assay

SNAPf–tagged Kinesin-8X-MD proteins (20 μM) were biotinylated in 50 μl reaction volumes by incubating with 40 μM SNAP-biotin (NEB) at 4 °C for 1.5 hrs. Proteins were purified from excess SNAP-biotin by size-exclusion chromatography on a Superdex 75 Increase 3.2/300 column using an ÅKTAmicro system (GE Healthcare) in gel filtration buffer (20 mM Tris-HCl pH 7.5, 250 mM NaCl, 5 mM MgCl_2_,1 mM DTT). Peak fractions were pooled, snap frozen in liquid nitrogen, and stored at −80°C until use. For MT gliding assays, flow chambers were treated and imaged as above except that in step 3) biotinylated Kinesin8X-MD was added instead of MTs and in step 4) the reaction mixture contained 5mM ATP together with 10 % X-rhodamine-MTs instead of Kinesin8X-MD proteins. Gliding assay movies were collected at room temperature for 10 min with 2 s interval. The gliding rates of single MTs were measured from kymographs using Fiji software.

### Ethics statement

The animal work performed in the UK passed an ethical review process and was approved by the United Kingdom Home Office. Work was carried out under UK Home Office Project Licenses (40/3344 and 30/3248) in accordance with the United Kingdom ‘Animals (Scientific Procedures) Act 1986’ and in compliance with ‘European Directive 86/609/EEC’ for the protection of animals used for experimental purposes. Experiments performed in Switzerland were conducted in strict accordance with the guidelines of the Swiss Tierschutzgesetz (TSchG; Animal Rights Laws) and approved by the ethical committee of the University of Bern (Permit Number: BE109/13). Six-to-eight week old female Tuck-Ordinary (TO) (Harlan) outbred mice were used for all experiments in the UK. Balb/c female mice between six and ten weeks of age were used in experiments in Switzerland. Mice were either bred in the central animal facility of the University of Bern, or were supplied by Harlan Laboratories or Charles River Laboratories.

### Generation of transgenic parasites

The C-terminus of Kinesin-8X was tagged with GFP by single crossover homologous recombination in the parasite. To generate the Kinesin-8X-GFP line, a region of the *kinesin-8* gene downstream of the ATG start codon was amplified using primers T1931 and T1932, ligated to p277 vector, and transfected as described previously [67]. A schematic representation of the endogenous *kinesin-8X* locus (PBANKA_080590), the constructs and the recombined *kinesin-8X* locus can be found in S2 Fig. The gene-deletion targeting vector for *kinesin-8X was* constructed using the pBS-DHFR plasmid, which contains polylinker sites flanking a *T. gondii dhfr/ts* expression cassette conferring resistance to pyrimethamine, as described previously [58]. PCR primers N1051 and N1052 were used to generate an 830 bp fragment of *kinesin-8X* 5′ upstream sequence from genomic DNA, which was inserted into *Apa*I and *Hin*dIII restriction sites upstream of the *dhfr/ts* cassette of pBS-DHFR. A 933 bp fragment generated with primers N1053 and N1054 from the 3′ flanking region of *kinesin-8X* was then inserted downstream of the *dhfr/ts* cassette using *Eco*RI and *Xba*I restriction sites. The linear targeting sequence was released using *Apa*I/*Xba*I. A schematic representation of the endogenous *kinesin-8X* locus (PBANKA_080590), the constructs and the recombined *kinesin-8X* locus can be found in S3 Fig. The oligonucleotides used to generate the mutant parasite lines can be found in S2 Table. *P. berghei* ANKA line 2.34 (for GFP-tagging) or ANKA line 507cl1 expressing GFP (for gene deletion) were transfected by electroporation and selected by pyrimethamine drug against di-hydro folate reductase selectable marker as described previously [68].

### Parasite genotype analyses

For the parasites expressing a C-terminal GFP-tagged Kinesin-8X protein, diagnostic PCR was used with primer 1 (IntT193) and primer 2 (ol492) to confirm integration of the GFP targeting construct. For the gene knockout parasites, diagnostic PCR was used with primer 1 (IntN105) and primer 2 (ol248) to confirm integration of the targeting construct, and primer 3 (N105 KO1) and primer 4 (N105 KO2) were used to confirm deletion of the *kinesin-8X* gene.

### Parasite phenotype analyses

Blood containing approximately 50,000 parasites of the kinesin-8X-KO line was injected intraperitoneally (i.p) into mice to initiate infections. Asexual stages and gametocyte production were monitored by microscopy on Giemsa stained thin smears. Four to five days post infection, exflagellation and ookinete conversion were examined as described previously [67] with a Zeiss AxioImager M2 microscope (Carl Zeiss, Inc) fitted with an AxioCam ICc1 digital camera. To analyse mosquito transmission, 30–50 *Anopheles stephensi* SD 500 mosquitoes were allowed to feed for 20 min on anaesthetized, infected mice whose asexual parasitaemia had reached 15% and were carrying comparable numbers of gametocytes as determined on Giemsa stained blood films. To assess mid-gut infection, approximately 15 guts were dissected from mosquitoes on day 14 post feeding and oocysts were counted on an AxioCam ICc1 digital camera fitted to a Zeiss AxioImager M2 microscope using a 63x oil immersion objective. On day 21 post-feeding, another 20 mosquitoes were dissected, and their guts and salivary glands crushed separately in a loosely fitting homogenizer to release sporozoites, which were then quantified using a haemocytometer or used for imaging. Mosquito bite back experiments were performed 21 days post-feeding using naive mice and blood smears were examined after 3-4 days.

### Electron microscopy

Mosquito midguts at 14-day post infection were fixed in 4% glutaraldehyde in 0.1 M phosphate buffer and processed for routine electron microscopy as previously described (56). Briefly, samples were post fixed in osmium tetroxide, treated en bloc with uranyl acetate, dehydrated and embedded in Spurr’s epoxy resin. Thin sections were stained with uranyl acetate and lead citrate prior to examination in a JEOL1200EX electron microscope (Jeol UK Ltd).

### Culture and gradient purification of schizonts and gametocytes

Blood cells obtained from infected mice (days 4 to 5 post infection) were placed in culture for 24 h at 37°C (with rotation at 100 rpm) and schizonts were purified the following day on a 60% v/v NycoDenz (in PBS) gradient, harvested from the interface and washed (NycoDenz stock solution: 27.6% w/v NycoDenz in 5 mM Tris-HCl, pH 7.20, 3 mM KCl, 0.3 mM EDTA). Purification of gametocytes was achieved using a protocol described as previously [69] with some modifications[70].

### Liver stage parasite imaging

For P. berghei liver stage parasites, 100,000 HeLa cells were seeded in glass-bottomed imaging dishes. Salivary glands of female *A. stephensi* mosquitoes infected with Kinesin-8X-GFP parasites were isolated and disrupted using a pestle to release sporozoites, which were pipetted gently onto the seeded HeLa cells and incubated at 37 °C in 5% CO_2_ in complete minimum Eagle’s medium containing 2.5 μg/ml amphotericin B (PAA). Medium was changed 3 hrs after initial infection and once a day thereafter. For live cell imaging, Hoechst 33342 (Molecular Probes) was added to a final concentration of 1 μg/ml, and parasites were imaged at 24, 48, 55 hrs post-infection using a Leica TCS SP8 confocal microscope with the HC PL APO 63x/1.40 oil objective and the Leica Application Suite X software

### Fixed Immunofluorescence Assay and DNA content analysis

The PbKinesin-8X-GFP gametocytes were purified and activated in ookinete medium then fixed at 0 min, 1-2 min, 6-8 min and 15 min post-activation with 4% paraformaldehyde (PFA, Sigma) diluted in microtubule stabilising buffer (MTSB) for 10-15 min and added to poly-L-lysine coated slides. Immunocytochemistry was performed using primary GFP-specific rabbit monoclonal antibody (mAb) (Invitrogen-A1122; used at 1:250) and primary mouse anti-α tubulin mAb (Sigma-T9026; used at 1:1000). Secondary antibodies were Alexa 488 conjugated anti-mouse IgG (Invitrogen-A11004) and Alexa 568 conjugated anti-rabbit IgG (Invitrogen-A11034) (used at 1 in 1000). The slides were then mounted in Vectashield 19 with DAPI (Vector Labs) for fluorescence microscopy. Parasites were visualised on a Zeiss AxioImager M2 microscope fitted with an AxioCam ICc1 digital camera (Carl Zeiss, Inc).

### Deconvolution microscopy

High resolution imaging was performed using an AxioCam ICc1 digital camera fitted to a Zeiss AxioImager M2 microscope using a 63x oil immersion objective. Post-acquisition analysis was carried out using Icy software - version 1.9.10.0. Images presented are 2D projections of deconvoluted Z-stacks of 0.3 μm optical sections.

### Ookinete motility assay and DNA content analysis

The assay was performed using matrigel as described previously (Volkmann et al., 2012) with some modification. Ookinete cultures were added to an equal volume of Matrigel (Corning) on ice, mixed thoroughly, dropped onto a slide, covered with a cover slip, and sealed with nail polish. The Matrigel was then allowed to set at 20°C for 30 minutes. After identifying a field containing ookinetes, time-lapse videos were taken at every 5 sec for 150 cycle.

DNA content of ookinete was analysed by Fluorometry after Hoechst nuclear staining as described previously (Guttery et al, 2014).

### Quantitative Real Time PCR (qRT-PCR) analyses

RNA was isolated from different parasite stages including asexual, purified schizonts, gametocytes, ookinete and sporozoites using an RNA purification kit (Stratagene). cDNA was synthesised using an RNA-to-cDNA kit (Applied Biosystems). Gene expression was quantified from 80 ng of total RNA using SYBR green fast master mix kit (Applied Biosystems). All the primers were designed using primer3 (Primer-blast, NCBI). Analysis was conducted using an Applied Biosystems 7500 fast machine with the following cycling conditions: 95°C for 20 s followed by 40 cycles of 95°C for 3 s; 60°C for 30 s. Three technical replicates and three biological replicates were performed for each assayed gene. The *hsp70* (PBANKA_081890) and *arginyl-t RNA synthetase* (PBANKA_143420) genes were used as endogenous control reference genes. The primers used for qPCR can be found in S2 Table.

### RNA-seq analysis

Libraries were prepared from lyophilized total RNA using the KAPA Library Preparation Kit (KAPA Biosystems). Libraries were amplified for a total of 12 PCR cycles (12 cycles of [15 s at 98°C, 30 s at 55°C, 30 s at 62°C]) using the KAPA HiFi HotStart Ready Mix (KAPA Biosystems). Libraries were sequenced using a NextSeq500 DNA sequencer (Illumina), producing paired-end 75-bp reads.

FastQC (https://www.bioinformatics.babraham.ac.uk/projects/fastqc/), was used to analyze raw read quality, and based on this information, the first 11 bp of each read and any adapter sequences were removed using Trimmomatic (http://www.usadellab.org/cms/?page=trimmomatic). Bases with Phred quality scores below 25 were trimmed using Sickle (https://github.com/najoshi/sickle). The resulting reads were mapped against the *Plasmodium berghei* ANKA genome (v36) using HISAT2 (version 2-2.1.0), using default parameters. Uniquely mapped, properly paired reads were retained using SAMtools (http://samtools.sourceforge.net/), and PCR duplicates were removed by PicardTools MarkDuplicates (Broad Institute). Genome browser tracks were generated and viewed using the Integrative Genomic Viewer (IGV) (Broad Institute).

Raw read counts were determined for each gene in the *P. berghei* genome using BedTools (https://bedtools.readthedocs.io/en/latest/#) to intersect the aligned reads with the genome annotation. Differential expression analysis was done by use of R package DESeq2 to call up- and down-regulated genes. Gene ontology enrichment was done on PlasmoDB (http://plasmodb.org/plasmo/) with repetitive terms removed by REVIGO (http://revigo.irb.hr/).

### Statistical analysis

All statistical analyses were performed using GraphPad Prism 7 (GraphPad Software). For qRT-PCR, an unpaired t-test was used to examine significant differences between wild-type and mutant strains.

### Data Availability

Sequence reads have been deposited in the NCBI Sequence Read Archive with accession number: <u>SUB5218526</u>

## Acknowledgments

We wish to thank Dr Michael Delves for advice on deconvolution microscopy, Robert E. Sinden for his valuable discussion and to Julie Rodgers for helping to maintain the insectary and other technical works.

## Funding

This work was supported by: MRC UK (G0900109, G0900278, MR/K011782/1) and BBSRC (BB/N017609/1) to RT; the BBSRC (BB/N017609/1) to MZ, TY and CAM; the EMBO-Long Term fellowship (597-2014) to MR; the Francis Crick Institute, the Cancer Research UK (FC001097), the UK Medical Research Council (FC001097), and the Wellcome Trust (FC001097) to AAH; the NIH/NIAID (R01 AI136511) and the University of California, Riverside (NIFA-Hatch-225935) to KGLR; a PhD studentship to F.S., a Sir Henry Dale Fellowship from the Wellcome Trust and Royal Society (104196/Z/14/Z) to AJR. Part of the workdone in AP laboratory was supported by the faculty baseline funding and a CRG3 grant from King Abdullah University of Science and Technology (KAUST).

---

**S1 Fig:**
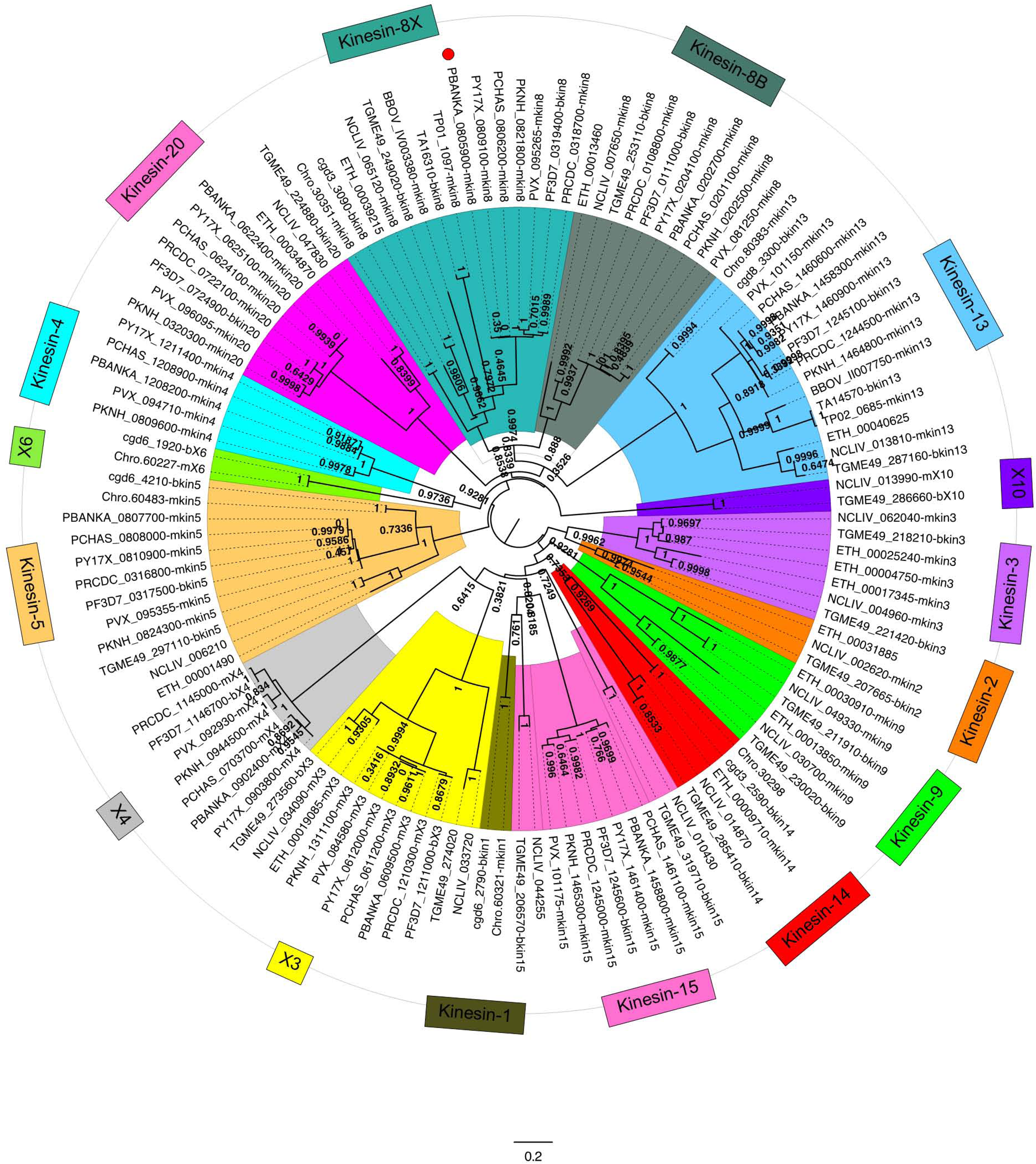
Phylogenetic analysis of kinesins in Apicomplexan. Phylogeny of detected kinesin protein sequences. Proteins with the suffix “-b[NNN]” are retrieved directly from Wickstead et al. [3], where NNN denote Kinesin gene. Proteins with the suffix “-m[NNN]” were also detected by the reciprocal best BLAST approach (see Methods). Tree produced using PhyML [71] with the LG+G+I+F model selected by SMS [72]. Branch support was evaluated with the Bayesian-like transformation of approximate likelihood ratio test (aBayes). Genetic distance shown below tree. Note that Kinesin-15 is a paraphyletic group.

**S2 Fig:**
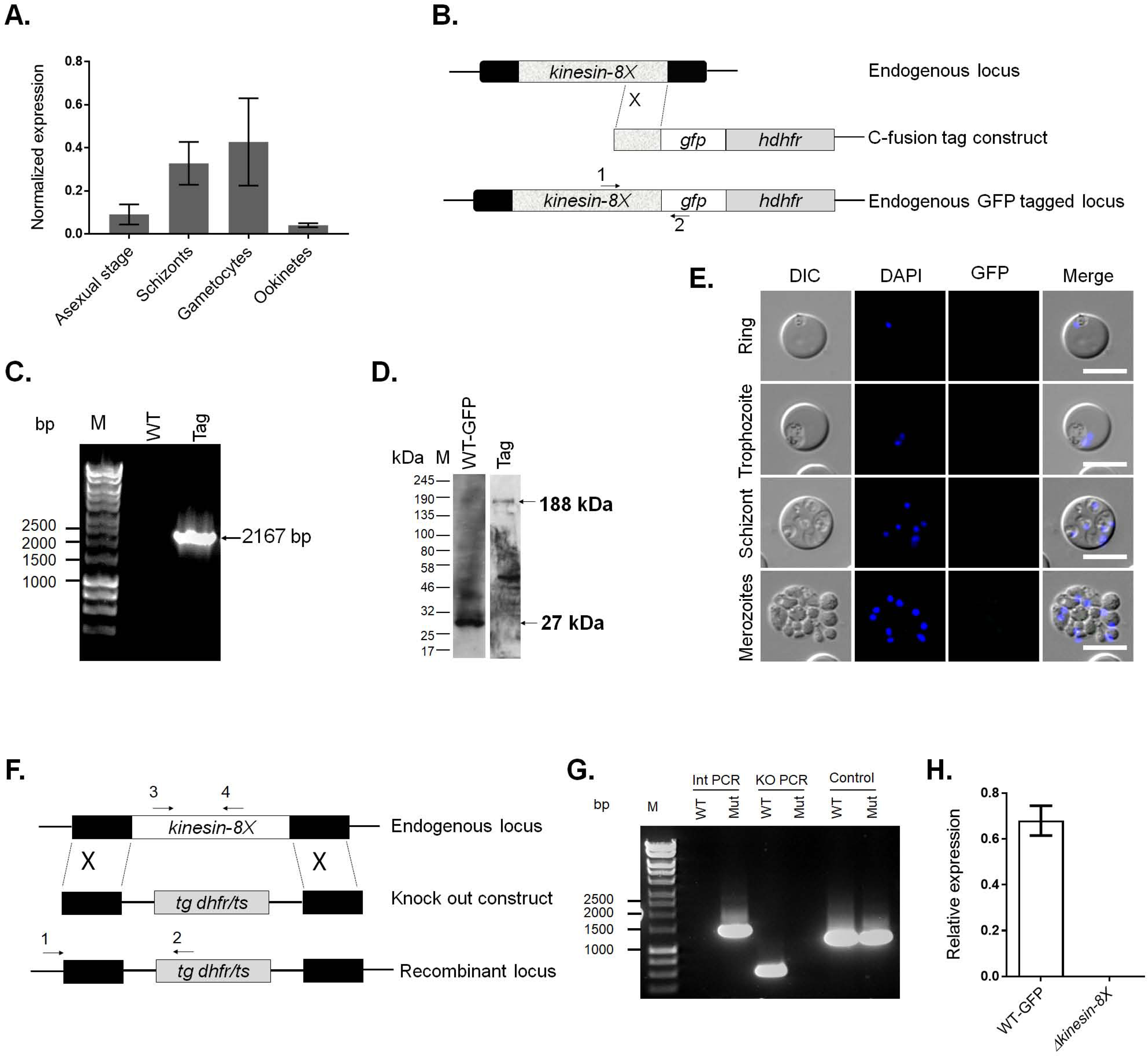
Generation and genotypic analysis of Kinesin-8X-GFP and *Δkinesin-8X* parasites. (A) Analysis of Kinesin-8X transcript level by qRT-PCR during different stages of *P. berghei* life cycle. Mean ± SD. n= 3 independent experiments. (B) Schematic representation of the endogenous *pbkinesin-8X* locus, the GFP-tagging construct and the recombined *kinesin-8X* locus following single homologous recombination. Arrows 1 and 2 indicate the position of PCR primers used to confirm successful integration of the construct. (C) Diagnostic PCR of *kinesin-8X* and WT parasites using primers IntT193 (Arrow 1) and ol492 (Arrow 2). Integration of the Kinesin-8X tagging construct gives a band of 2167 bp. Tag = kinesin-8X-GFP parasite line. (D) Western blot of Kinesin-8X-GFP (~188 kDa) and WT-GFP (~27 kDa) protein to illustrate Kinesin-8X-GFP in gametocyte stage. (E) Live cell imaging of Kinesin-8X-GFP parasites during erythrocytic schizogony (F) Schematic representation of the endogenous kinesin-8x locus, the targeting knockout construct and the recombined kinesin-8X locus following double homologous cross-over recombination. Arrows 1 and 2 indicate PCR primers used to confirm successful integration in the kinesin-8X locus following recombination and arrows 3 and 4 indicate PCR primers used to show deletion of the kinesin-8X gene. (G) Integration PCR of the kinesin-8X locus in WT-GFP and *Δkinesin-8X* (Mut) parasites using primers INT N105 and ol248. Integration of the targeting construct gives a band of 1.5 kb. (H) qRT-PCR analysis of transcript in WT-GFP and *Δkinesin-8X* parasites. Mean ± SD. n= 3 independent experiments.

**S3 Fig:** Location of Kinesin-8X with kinetochore marker Ndc80. Live cell imaging showing that Kinesin-8X is located next to Ndc80, a kinetochore marker, in oocysts stage, suggesting that it is not part of kinetochore. It is clearer in sporozoites where kinesin-8X is enriched next to nucleus and Ndc80.

**S4 Fig:** Analysis of morphology, DNA content and motility of *Δkinesin-8X* ookinetes. (A) Morphology of ookinetes showing no difference in WT-GFP and *Δkinesin-8X* parasites. (B) Fluorometric DNA content (N) analysis of WT-GFP and *Δkinesin-8X* ookinetes, after Hoechst nuclear staining. Nuclear fluorescence intensity of WT-GFP or mutant parasites from 24 hr cultures was measured using ImageJ software. Values are expressed relative to the average fluorescence intensity of haploid ring-stage parasites from the same slide and corrected for background fluorescence (Error bar ±SD; n=3 independent experiments, >10 ookinetes were analysed for each experiment).

**S1 Table. Phylogenetic analysis of Kinesins in Apicomplexans**

**S2 Table. Oligonucleotides used in this study**

**S3 Table. List of differentially expressed genes between Δ*kinesin-8X* and WT activated gametocytes**

## Notes

#### Summary of Updates

1. Small changes in abstract and author summary 2. Some updates in the method section 3. Moving the name Robert E. Sinden from author list to acknowledgement due to conflict of interest 4. Few updates in legends

## References

1. Lawrence CJ, Dawe RK, Christie KR, Cleveland DW, Dawson SC, Endow SA, et al. A standardized kinesin nomenclature. J Cell Biol. 2004;167(1):19–22. doi: 10.1083/jcb.200408113. PubMed PMID: 15479732; PubMed Central PMCID: PMCPMC2041940.

2. Vicente JJ, Wordeman L. Mitosis, microtubule dynamics and the evolution of kinesins. Exp Cell Res. 2015;334(1):61–9. doi: 10.1016/j.yexcr.2015.02.010. PubMed PMID: 25708751; PubMed Central PMCID: PMCPMC4433793.

3. Wickstead B, Gull K, Richards TA. Patterns of kinesin evolution reveal a complex ancestral eukaryote with a multifunctional cytoskeleton. BMC Evol Biol. 2010;10:110. PubMed PMID: 20423470.

4. Verhey KJ, Hammond JW. Traffic control: regulation of kinesin motors. Nat Rev Mol Cell Biol. 2009;10(11):765–77. PubMed PMID: 19851335.

5. Prosser SL, Pelletier L. Mitotic spindle assembly in animal cells: a fine balancing act. Nat Rev Mol Cell Biol. 2017;18(3):187–201. doi: 10.1038/nrm.2016.162. PubMed PMID: 28174430.

6. Wittmann T, Hyman A, Desai A. The spindle: a dynamic assembly of microtubules and motors. Nat Cell Biol. 2001;3(1):E28–34. doi: 10.1038/35050669. PubMed PMID: 11146647.

7. Kushida Y, Takaine M, Nakano K, Sugai T, Vasudevan KK, Guha M, et al. Kinesin-14 is Important for Chromosome Segregation During Mitosis and Meiosis in the Ciliate Tetrahymena thermophila. J Eukaryot Microbiol. 2017;64(3):293–307. doi: 10.1111/jeu.12366. PubMed PMID: 27595611.

8. Bascom-Slack CA, Dawson DS. The yeast motor protein, Kar3p, is essential for meiosis I. J Cell Biol. 1997;139(2):459–67. PubMed PMID: 9334348; PubMed Central PMCID: PMCPMC2139793.

9. Camlin NJ, McLaughlin EA, Holt JE. Motoring through: the role of kinesin superfamily proteins in female meiosis. Hum Reprod Update. 2017;23(4):409–20. doi: 10.1093/humupd/dmx010. PubMed PMID: 28431155.

10. Mayr MI, Hummer S, Bormann J, Gruner T, Adio S, Woehlke G, et al. The human kinesin Kif18A is a motile microtubule depolymerase essential for chromosome congression. Curr Biol. 2007;17(6):488–98. doi: 10.1016/j.cub.2007.02.036. PubMed PMID: 17346968.

11. DeZwaan TM, Ellingson E, Pellman D, Roof DM. Kinesin-related KIP3 of Saccharomyces cerevisiae is required for a distinct step in nuclear migration. J Cell Biol. 1997;138(5):1023–40. PubMed PMID: 9281581; PubMed Central PMCID: PMCPMC2136764.

12. Stumpff J, von Dassow G, Wagenbach M, Asbury C, Wordeman L. The kinesin-8 motor Kif18A suppresses kinetochore movements to control mitotic chromosome alignment. Dev Cell. 2008;14(2):252–62. doi: 10.1016/j.devcel.2007.11.014. PubMed PMID: 18267093; PubMed Central PMCID: PMCPMC2267861.

13. Savoian MS, Gatt MK, Riparbelli MG, Callaini G, Glover DM. Drosophila Klp67A is required for proper chromosome congression and segregation during meiosis I. J Cell Sci. 2004;117(Pt 16):3669–77. PubMed PMID: 15252134.

14. Mary H, Fouchard J, Gay G, Reyes C, Gauthier T, Gruget C, et al. Fission yeast kinesin-8 controls chromosome congression independently of oscillations. J Cell Sci. 2015;128(20):3720–30. PubMed PMID: 26359299.

15. Straight AF, Sedat JW, Murray AW. Time-lapse microscopy reveals unique roles for kinesins during anaphase in budding yeast. J Cell Biol. 1998;143(3):687–94. PubMed PMID: 9813090; PubMed Central PMCID: PMCPMC2148141.

16. Savoian MS, Glover DM. Drosophila Klp67A binds prophase kinetochores to subsequently regulate congression and spindle length. J Cell Sci. 2010;123(Pt 5):767–76. doi: 10.1242/jcs.055905. PubMed PMID: 20144994.

17. Gupta ML, Jr., Carvalho P, Roof DM, Pellman D. Plus end-specific depolymerase activity of Kip3, a kinesin-8 protein, explains its role in positioning the yeast mitotic spindle. Nat Cell Biol. 2006;8(9):913–23. doi: 10.1038/ncb1457. PubMed PMID: 16906148.

18. Varga V, Helenius J, Tanaka K, Hyman AA, Tanaka TU, Howard J. Yeast kinesin-8 depolymerizes microtubules in a length-dependent manner. Nat Cell Biol. 2006;8(9):957–62. PubMed PMID: 16906145.

19. Wang D, Nitta R, Morikawa M, Yajima H, Inoue S, Shigematsu H, et al. Motility and microtubule depolymerization mechanisms of the Kinesin-8 motor, KIF19A. Elife. 2016;5. PubMed PMID: 27690357.

20. Tran PT, Doye V, Chang F, Inoue S. Microtubule-dependent nuclear positioning and nuclear-dependent septum positioning in the fission yeast Schizosaccharomyces [correction of Saccharomyces] pombe. Biol Bull. 2000;199(2):205–6. doi: 10.2307/1542900. PubMed PMID: 11081738.

21. West RR, Malmstrom T, Troxell CL, McIntosh JR. Two related kinesins, klp5+ and klp6+, foster microtubule disassembly and are required for meiosis in fission yeast. Mol Biol Cell. 2001;12(12):3919–32. doi: 10.1091/mbc.12.12.3919. PubMed PMID: 11739790; PubMed Central PMCID: PMCPMC60765.

22. Daga RR, Yonetani A, Chang F. Asymmetric microtubule pushing forces in nuclear centering. Curr Biol. 2006;16(15):1544–50. doi: 10.1016/j.cub.2006.06.026. PubMed PMID: 16890530.

23. WHO. World Malaria Report. 2018.

24. Sinden RE. Mitosis and meiosis in malarial parasites. Acta Leiden. 1991;60(1):19–27. PubMed PMID: 1820709.

25. Francia ME, Striepen B. Cell division in apicomplexan parasites. Nat Rev Microbiol. 2014;12(2):125–36. PubMed PMID: 24384598.

26. Arnot DE, Ronander E, Bengtsson DC. The progression of the intra-erythrocytic cell cycle of Plasmodium falciparum and the role of the centriolar plaques in asynchronous mitotic division during schizogony. Int J Parasitol. 2011;41(1):71–80. PubMed PMID: 20816844.

27. Sinden RE. Sexual development of malarial parasites. Adv Parasitol. 1983;22:153–216. PubMed PMID: 6141715.

28. Billker O, Shaw MK, Margos G, Sinden RE. The roles of temperature, pH and mosquito factors as triggers of male and female gametogenesis of Plasmodium berghei in vitro. Parasitology. 1997;115 (Pt 1):1–7. PubMed PMID: 9280891.

29. Sinden RE, Canning EU, Bray RS, Smalley ME. Gametocyte and gamete development in Plasmodium falciparum. Proc R Soc Lond B Biol Sci. 1978;201(1145):375–99. PubMed PMID: 27809.

30. Guttery DS, Roques M, Holder AA, Tewari R. Commit and Transmit: Molecular Players in Plasmodium Sexual Development and Zygote Differentiation. Trends Parasitol. 2015;31(12):676–85. doi: 10.1016/j.pt.2015.08.002. PubMed PMID: 26440790.

31. Schrevel J, Asfaux-Foucher G, Bafort JM. [Ultrastructural study of multiple mitoses during sporogony of Plasmodium b. berghei]. J Ultrastruct Res. 1977;59(3):332–50. PubMed PMID: 864828.

32. Sinden RE. Gametocytogenesis of Plasmodium falciparum in vitro: an electron microscopic study. Parasitology. 1982;84(1):1–11. PubMed PMID: 7038594.

33. Roques M, Stanway RR, Rea EI, Markus R, Brady D, Holder AA, et al. Plasmodium centrin PbCEN-4 localizes to the putative MTOC and is dispensable for malaria parasite proliferation. Biol Open. 2019;8(1). doi: 10.1242/bio.036822. PubMed PMID: 30541825.

34. Gerald N, Mahajan B, Kumar S. Mitosis in the human malaria parasite Plasmodium falciparum. Eukaryot Cell. 2011;10(4):474–82. doi: 10.1128/EC.00314-10. PubMed PMID: 21317311; PubMed Central PMCID: PMCPMC3127633.

35. Wandke C, Barisic M, Sigl R, Rauch V, Wolf F, Amaro AC, et al. Human chromokinesins promote chromosome congression and spindle microtubule dynamics during mitosis. J Cell Biol. 2012;198(5):847–63. doi: 10.1083/jcb.201110060. PubMed PMID: 22945934; PubMed Central PMCID: PMCPMC3432768.

36. Cross RA, McAinsh A. Prime movers: the mechanochemistry of mitotic kinesins. Nat Rev Mol Cell Biol. 2014;15(4):257–71. doi: 10.1038/nrm3768. PubMed PMID: 24651543.

37. Liu L, Richard J, Kim S, Wojcik EJ. Small molecule screen for candidate antimalarials targeting Plasmodium Kinesin-5. J Biol Chem. 2014;289(23):16601–14. doi: 10.1074/jbc.M114.551408. PubMed PMID: 24737313; PubMed Central PMCID: PMCPMC4047425.

38. Locke J, Joseph AP, Pena A, Mockel MM, Mayer TU, Topf M, et al. Structural basis of human kinesin-8 function and inhibition. Proc Natl Acad Sci U S A. 2017;114(45):E9539–E48. doi: 10.1073/pnas.1712169114. PubMed PMID: 29078367; PubMed Central PMCID: PMCPMC5692573.

39. Otto TD, Bohme U, Jackson AP, Hunt M, Franke-Fayard B, Hoeijmakers WA, et al. A comprehensive evaluation of rodent malaria parasite genomes and gene expression. BMC Biol. 2014;12:86. doi: 10.1186/s12915-014-0086-0. PubMed PMID: 25359557; PubMed Central PMCID: PMCPMC4242472.

40. Yeoh LM, Goodman CD, Mollard V, McFadden GI, Ralph SA. Comparative transcriptomics of female and male gametocytes in Plasmodium berghei and the evolution of sex in alveolates. BMC Genomics. 2017;18(1):734. doi: 10.1186/s12864-017-4100-0. PubMed PMID: 28923023; PubMed Central PMCID: PMCPMC5604118.

41. Billker O, Lindo V, Panico M, Etienne AE, Paxton T, Dell A, et al. Identification of xanthurenic acid as the putative inducer of malaria development in the mosquito. Nature. 1998;392(6673):289–92. doi: 10.1038/32667. PubMed PMID: 9521324.

42. Janse CJ, Mons B, Rouwenhorst RJ, Van der Klooster PF, Overdulve JP, Van der Kaay HJ. In vitro formation of ookinetes and functional maturity of Plasmodium berghei gametocytes. Parasitology. 1985;91 (Pt 1):19–29. PubMed PMID: 2863802.

43. Bushell E, Gomes AR, Sanderson T, Anar B, Girling G, Herd C, et al. Functional Profiling of a Plasmodium Genome Reveals an Abundance of Essential Genes. Cell. 2017;170(2):260–72 e8. doi: 10.1016/j.cell.2017.06.030. PubMed PMID: 28708996; PubMed Central PMCID: PMCPMC5509546.

44. Messin LJ, Millar JB. Role and regulation of kinesin-8 motors through the cell cycle. Syst Synth Biol. 2014;8(3):205–13. doi: 10.1007/s11693-014-9140-z. PubMed PMID: 25136382; PubMed Central PMCID: PMCPMC4127180.

45. Su X, Arellano-Santoyo H, Portran D, Gaillard J, Vantard M, Thery M, et al. Microtubule-sliding activity of a kinesin-8 promotes spindle assembly and spindle-length control. Nat Cell Biol. 2013;15(8):948–57. doi: 10.1038/ncb2801. PubMed PMID: 23851487; PubMed Central PMCID: PMCPMC3767134.

46. Shrestha S, Hazelbaker M, Yount AL, Walczak CE. Emerging Insights into the Function of Kinesin-8 Proteins in Microtubule Length Regulation. Biomolecules. 2018;9(1). doi: 10.3390/biom9010001. PubMed PMID: 30577528; PubMed Central PMCID: PMCPMC6359247.

47. Gergely ZR, Crapo A, Hough LE, McIntosh JR, Betterton MD. Kinesin-8 effects on mitotic microtubule dynamics contribute to spindle function in fission yeast. Mol Biol Cell. 2016;27(22):3490–514. doi: 10.1091/mbc.E15-07-0505. PubMed PMID: 27146110; PubMed Central PMCID: PMCPMC5221583.

48. Unsworth A, Masuda H, Dhut S, Toda T. Fission yeast kinesin-8 Klp5 and Klp6 are interdependent for mitotic nuclear retention and required for proper microtubule dynamics. Mol Biol Cell. 2008;19(12):5104–15. doi: 10.1091/mbc.E08-02-0224. PubMed PMID: 18799626; PubMed Central PMCID: PMCPMC2592636.

49. Tytell JD, Sorger PK. Analysis of kinesin motor function at budding yeast kinetochores. J Cell Biol. 2006;172(6):861–74. doi: 10.1083/jcb.200509101. PubMed PMID: 16533946; PubMed Central PMCID: PMCPMC2063730.

50. Goshima G, Vale RD. Cell cycle-dependent dynamics and regulation of mitotic kinesins in Drosophila S2 cells. Mol Biol Cell. 2005;16(8):3896–907. doi: 10.1091/mbc.e05-02-0118. PubMed PMID: 15958489; PubMed Central PMCID: PMCPMC1182325.

51. Sinden RE, Hartley RH. Identification of the meiotic division of malarial parasites. J Protozool. 1985;32(4):742–4. PubMed PMID: 3906103.

52. Canning EU, Sinden RE. The organization of the ookinete and observations on nuclear division in oocysts of Plasmodium berghei. Parasitology. 1973;67(1):29–40. PubMed PMID: 4579580.

53. Sinden RE, Strong K. An ultrastructural study of the sporogonic development of Plasmodium falciparum in Anopheles gambiae. Trans R Soc Trop Med Hyg. 1978;72(5):477–91. PubMed PMID: 364785.

54. Walczak CE, Mitchison TJ, Desai A. XKCM1: a Xenopus kinesin-related protein that regulates microtubule dynamics during mitotic spindle assembly. Cell. 1996;84(1):37–47. PubMed PMID: 8548824.

55. Moores CA, Yu M, Guo J, Beraud C, Sakowicz R, Milligan RA. A mechanism for microtubule depolymerization by KinI kinesins. Mol Cell. 2002;9(4):903–9. PubMed PMID: 11983180.

56. s M, Wall RJ, Douglass AP, Ramaprasad A, Ferguson DJ, Kaindama ML, et al. Plasmodium P-Type Cyclin CYC3 Modulates Endomitotic Growth during Oocyst Development in Mosquitoes. PLoS Pathog. 2015;11(11):e1005273. doi: 10.1371/journal.ppat.1005273. PubMed PMID: 26565797; PubMed Central PMCID: PMCPMC4643991.

57. Bushell ES, Ecker A, Schlegelmilch T, Goulding D, Dougan G, Sinden RE, et al. Paternal effect of the nuclear formin-like protein MISFIT on Plasmodium development in the mosquito vector. PLoS Pathog. 2009;5(8):e1000539. doi: 10.1371/journal.ppat.1000539. PubMed PMID: 19662167; PubMed Central PMCID: PMCPMC2715856.

58. Tewari R, Straschil U, Bateman A, Bohme U, Cherevach I, Gong P, et al. The systematic functional analysis of Plasmodium protein kinases identifies essential regulators of mosquito transmission. Cell Host Microbe. 2010;8(4):377–87. doi: 10.1016/j.chom.2010.09.006. PubMed PMID: 20951971; PubMed Central PMCID: PMCPMC2977076.

59. Mlambo G, Coppens I, Kumar N. Aberrant sporogonic development of Dmc1 (a meiotic recombinase) deficient Plasmodium berghei parasites. PLoS One. 2012;7(12):e52480. doi: 10.1371/journal.pone.0052480. PubMed PMID: 23285059; PubMed Central PMCID: PMCPMC3528682.

60. Guttery DS, Poulin B, Ramaprasad A, Wall RJ, Ferguson DJ, Brady D, et al. Genome-wide functional analysis of Plasmodium protein phosphatases reveals key regulators of parasite development and differentiation. Cell Host Microbe. 2014;16(1):128–40. doi: 10.1016/j.chom.2014.05.020. PubMed PMID: 25011111; PubMed Central PMCID: PMCPMC4094981.

61. Deligianni E, Silmon de Monerri NC, McMillan PJ, Bertuccini L, Superti F, Manola M, et al. Essential role of Plasmodium perforin-like protein 4 in ookinete midgut passage. PLoS One. 2018;13(8):e0201651. doi: 10.1371/journal.pone.0201651. PubMed PMID: 30102727; PubMed Central PMCID: PMCPMC6089593.

62. Cottingham FR, Hoyt MA. Mitotic spindle positioning in Saccharomyces cerevisiae is accomplished by antagonistically acting microtubule motor proteins. J Cell Biol. 1997;138(5):1041–53. PubMed PMID: 9281582; PubMed Central PMCID: PMCPMC2136752.

63. El-Gebali S, Mistry J, Bateman A, Eddy SR, Luciani A, Potter SC, et al. The Pfam protein families database in 2019. Nucleic Acids Res. 2019;47(D1):D427–D32. doi: 10.1093/nar/gky995. PubMed PMID: 30357350; PubMed Central PMCID: PMCPMC6324024.

64. Katoh K, Standley DM. MAFFT multiple sequence alignment software version 7: improvements in performance and usability. Mol Biol Evol. 2013;30(4):772–80. doi: 10.1093/molbev/mst010. PubMed PMID: 23329690; PubMed Central PMCID: PMCPMC3603318.

65. Capella-Gutierrez S, Silla-Martinez JM, Gabaldon T. trimAl: a tool for automated alignment trimming in large-scale phylogenetic analyses. Bioinformatics. 2009;25(15):1972–3. doi: 10.1093/bioinformatics/btp348. PubMed PMID: 19505945; PubMed Central PMCID: PMCPMC2712344.

66. Hackney DD, Jiang W. Assays for kinesin microtubule-stimulated ATPase activity. Methods Mol Biol. 2001;164:65–71. PubMed PMID: 11217616.

67. Guttery DS, Poulin B, Ferguson DJ, Szoor B, Wickstead B, Carroll PL, et al. A unique protein phosphatase with kelch-like domains (PPKL) in Plasmodium modulates ookinete differentiation, motility and invasion. PLoS Pathog. 2012;8(9):e1002948. doi: 10.1371/journal.ppat.1002948. PubMed PMID: 23028336; PubMed Central PMCID: PMCPMC3447748.

68. Janse CJ, Ramesar J, Waters AP. High-efficiency transfection and drug selection of genetically transformed blood stages of the rodent malaria parasite Plasmodium berghei. Nat Protoc. 2006;1(1):346–56. doi: 10.1038/nprot.2006.53. PubMed PMID: 17406255.

69. Beetsma AL, van de Wiel TJ, Sauerwein RW, Eling WM. Plasmodium berghei ANKA: purification of large numbers of infectious gametocytes. Exp Parasitol. 1998;88(1):69–72. doi: 10.1006/expr.1998.4203. PubMed PMID: 9501851.

70. Saini E, Zeeshan M, Brady D, Pandey R, Kaiser G, Koreny L, et al. Photosensitized INA-Labelled protein 1 (PhIL1) is novel component of the inner membrane complex and is required for Plasmodium parasite development. Sci Rep. 2017;7(1):15577. doi: 10.1038/s41598-017-15781-z. PubMed PMID: 29138437; PubMed Central PMCID: PMCPMC5686188.

71. Guindon S, Dufayard JF, Lefort V, Anisimova M, Hordijk W, Gascuel O. New algorithms and methods to estimate maximum-likelihood phylogenies: assessing the performance of PhyML 3.0. Syst Biol. 2010;59(3):307–21. doi: 10.1093/sysbio/syq010. PubMed PMID: 20525638.

72. Lefort V, Longueville JE, Gascuel O. SMS: Smart Model Selection in PhyML. Mol Biol Evol. 2017;34(9):2422–4. doi: 10.1093/molbev/msx149. PubMed PMID: 28472384; PubMed Central PMCID: PMCPMC5850602.

